# Adipose tissue protein kinase D (PKD): regulation of signalling networks and its sex-dependent effects on metabolism

**DOI:** 10.1101/2024.08.29.610294

**Authors:** Mark C. Renton, Sean J. Humphrey, Tim Connor, Sheree D. Martin, Krystal Kremerer, Hilary Fernando, Christopher S. Shaw, David E. James, Kirsten F. Howlett, Sean L. McGee

## Abstract

The protein kinase D (PKD) family of three highly homologous isoforms (PKD1, PKD2, and PKD3) are implicated as nutrient sensing signalling kinases that regulate the response of adipose and other tissues to the nutrient environment. However, the physiological role of adipose tissue PKD and its downstream cellular signalling targets are not well characterised. Phosphoproteomics was performed to elucidate signalling events downstream of PKD activation in differentiated 3T3L1 adipocytes using a triple isoform siRNA knockdown model. This revealed PKD-regulated pathways including insulin and cAMP signalling, which control metabolic responses in adipose tissue. An adipose tissue-specific and inducible dominant negative PKD (atDNPKD) mouse model that achieves functional inhibition of all three PKD isoforms was generated to assess the function of adipose PKD on whole-body metabolism *in vivo* in both male and female mice. Insulin-stimulated suppression of lipolysis was blunted in male, but not female, atDNPKD mice compared to control mice. Female, but not male, atDNPKD mice had higher fasting insulin but normal insulin action. Male atDNPKD mice showed greater sensitivity to the β_3_-adrenergic receptor agonist CL316,243 on measures of lipolysis and energy expenditure, and displayed greater fat oxidation during fasting. During refeeding, male atDNPKD mice consumed less food and took longer to regain body weight lost during fasting. These effects were not observed in female mice. These findings indicate that PKD provides sex-dependent fine-tuning control of cAMP signalling in adipose tissue that is important for the coordination of energy balance during fasting and refeeding.

**NEW & NOTEWORTHY:** The protein kinase D (PKD) family is a target for the treatment of obesity-related disorders. However, the physiological role of PKD in adipose tissue remains to be resolved. Using phosphoproteomics and an adipose tissue PKD loss-of-function mouse model, results demonstrate that PKD provides fine tuning of metabolic signalling in adipose tissue and metabolic responses to fasting and refeeding challenges, via coordination of feeding behaviour and regulation of body weight.

## INTRODUCTION

Adipose tissue is the principal site for energy storage and is intimately involved in metabolic responses to nutrient availability. In the post prandial state, adipose tissue stores excess energy as triglycerides, a process coordinated by insulin (1). During fasting, catecholamines and other hormones liberate fatty acids from adipose tissue through lipolysis (2). Adipose tissue also secretes factors that provide information to other tissues on whole body energy balance (3). Understanding how these processes are coordinated at a molecular level has relevance for obesity, which is characterised by adipose tissue accumulation and dysfunction (4–6).

Protein kinase D (PKD) is a stress-activated regulatory kinase that controls a variety of essential cellular processes including cell differentiation and proliferation, inflammation, and exosomal release from the trans-Golgi network (7–9). The PKD family consists of three structurally similar isoforms (PKD1, PKD2 and PKD3) and share highly conserved C-terminal catalytic domains with homology to the CaMK kinase family (10). Canonical PKD activation, including the required phosphorylation sites, have been expertly reviewed previously (11, 12). In brief, G-protein-coupled receptor agonism results in phosphorylation of PKD at two activation loop serine sites (PKD1 Ser^744/748^) in a protein kinase C (PKC)-dependent pathway. This leads to subsequent autophosphorylation of PKD at its C-terminus (PKD1 Ser^916^), which is commonly used as a measure of PKD activity. Importantly, PKD activation also requires diacylglycerol and/or oxidative stress (13), both of which are upregulated in many tissues in obesity (14–18). As such, PKD has been postulated to be a key nutrient sensor in tissues important for systemic metabolism, such as the heart, liver, skeletal muscle, pancreas and adipose tissue, with potential implications for the development of obesity (11, 19). In adipose tissue, the activation pattern of PKD changes in response to feeding, with mice fasted for 24 hrs showing marked increases in PKD1 gene and protein expression. Furthermore, increases in PKD Ser^916^ phosphorylation in subcutaneous white adipose tissue are observed following refeeding (20). Adipose tissue-specific knockout of PKD1 in mice fed a high-fat diet (HFD) for 24 weeks resulted in significantly reduced weight gain, fat mass and liver steatosis, while energy expenditure, glucose tolerance, insulin sensitivity and markers of white adipose tissue beiging were all increased compared to control mice (20).

Although PKD activity could be involved in adipose tissue dysfunction in obesity, the biological functions regulated by adipose tissue PKD remain to be identified. In lean mice, adipose PKD1 knockout reduced adipocyte size in subcutaneous white adipose tissue and suppressed lipogenesis when compared to control littermates, suggesting an important role in metabolic coordination (21). Furthermore, there are currently very few identified and well-characterised targets of PKD within adipose tissue. PKD is known to inhibit 5’ adenosine monophosphate-activated protein kinase (AMPK) via inhibitory phosphorylation of Ser^485/491^ (22). AMPK activity was required for reduced lipogenesis in PKD1 knockout adipocytes (20), suggesting AMPK is a bona fide target of PKD1 in adipose tissue. PKD has well-established roles in the exocytosis of secretory vesicles from the Golgi (23), and evidence suggests that PKD2 is required for the insulin-stimulated secretion of lipoprotein lipase (LPL) from adipocytes via this mechanism (24).

A barrier to the study of PKD signalling is the highly homologous structure of the PKD family isoforms which results in functional redundancy in multiple cell types (24–27). For example, all three PKD isoforms phosphorylate histone deacetylase 5 (HDAC5) at Ser^498^ (26). Both PKD1 and PKD2 regulate LPL secretion from cardiomyocytes (27) and adipocytes (24), respectively. Additionally, genetic PKD1 knockdown causes a compensatory increase in PKD2 protein expression in cardiomyocytes (25). As such, care must be taken in interpreting data from single-isoform knockout experiments, as key signalling events or phenotypes can be compensated for by the remaining two isoforms. Despite this known redundancy, much of the literature investigating the role of PKD has focused on one isoform in isolation due to the difficulty of targeting multiple isoforms in a single experiment (20, 21, 24, 28).

There remains a gap in the literature describing the cellular signalling pathways that are regulated by the PKD family of isoforms within adipocytes. The current study employed a phosphoproteomics-based approach to map the downstream signalling targets of PKD in differentiated 3T3L1 adipocytes, and subsequently characterized a novel mouse model of PKD inactivation. Importantly, this study addresses isoform redundancy by designing both *in vitro* and *in* vivo models that genetically inhibit all three PKD isoforms. Results establish PKD as a regulator of key metabolic pathways in adipose tissue.

## MATERIALS AND METHODS

### Cell culture

3T3-L1 mouse fibroblasts (CL-173, ATCC, USA) were cultured in Dulbecco’s Modified Eagle Medium (DMEM; Gibco, USA) with 10% heat-inactivated fetal bovine serum (FBS; AusGeneX, Australia) and 1x penicillin-streptomycin (Gibco, USA) at 37°C in 10% CO_2_. To initiate differentiation into adipocytes, cells were grown until 3 days post-confluency and media was then changed to DMEM growth media supplemented with 0.4 µM insulin (Novo Nordisk, Denmark), 0.25 µM dexamethasone (Sigma-Aldrich, USA) and 500 µM isobutyl-1-methyl-xanthine (IBMX; Sigma-Aldrich, USA) for three days. Cells were then incubated in DMEM growth media supplemented only with 0.4 µM insulin for 4 days. Cells were then maintained in DMEM growth media without antibiotics refreshing media every 2-3 days.

To knockdown expression of PKD, 3T3-L1 adipocytes, at one-to-three days post-differentiation were transfected with siRNAs against Prkd1 (ON-TARGETplus Mouse Prkd1 siRNA SMARTPool; L-048415-00-0005; Dharmacon, USA), Prkd2 (ON-TARGETplus Mouse Prkd2 siRNA SMARTPool; L-040693-00-0005; Dharmacon, USA) and Prkd3 (ON-TARGETplus Mouse Prkd2 siRNA SMARTPool; L-040692-00-0005; Dharmacon, USA), or a non-targeting control pool siRNA (scramble siRNA; ON-TARGETplus Non-targeting pool; D-001810-10-20; Dharmacon, USA). Prkd siRNAs were transfected individually, or together (siPKD1/2/3) as indicated in individual experiments. The transfection protocol was adapted from a previously published method (29). Prior to transfection, RNAiMAX (Invitrogen, USA) was diluted in Opti-MEM reduced serum medium (Gibco, USA). Individual siRNA was also diluted in Opti-MEM before diluted RNAiMAX and siRNA mixes were combined and incubated at room temperature for 25 min. Combined RNAiMAX and siRNA mix was added to cells at 200 nM (optimised in test experiments) in antibiotic-free DMEM growth media and incubated for 4 days, when experiments were completed.

Scramble and siPKD1/2/3 transfected 3T3-L1 differentiated adipocytes were serum-starved in non-supplemented DMEM media for 30 min. Cells were then treated with phorbol 12-myristate 13-acetate (PMA), to activate PKD (25), at 5 μM (optimised in test experiments) or an equal volume of dimethyl sulfoxide (DMSO) as a vehicle control. Following PMA or vehicle treatment, adipocytes were washed in PBS and then lysed in ice-cold protein lysis buffer (1% Triton X-100, 10% glycerol, 1 mM Ethylenediaminetetraacetic acid, 1 mM ethylene glycol-bis(β-aminoethyl ether)-N,N,Nʹ,Nʹ-tetraacetic acid, 50 mM sodium fluoride, 5 mM sodium pyrophosphate, 1 mM sodium orthovanadate, 1 mM dithiothreitol, 50 mM Tris HCl pH 7.5 and 0.2% protease inhibitor cocktail (P8340; Sigma, USA) in Milli-Q water) via mechanical scraping. Cell lysate was then sonicated (2× 10 s pulses) using a tip probe sonicator (Qsonica, USA) and rotated at a medium speed on a rotating suspension mixer (Ratek, Australia) at 4°C for 2 hrs. Cell lysates were then spun in a centrifuge at 16 500 g for 15 min at 4°C to separate lipids from the protein homogenate. The purified protein homogenate was carefully removed without disturbing the pellet or top lipid layer, placed into a separate tube, and stored at –80°C for further analysis as described below. An aliquot of protein was diluted 1:5 in Milli-Q water and protein concentration was quantified using a Pierce™ BCA Protein Assay kit (Thermo Scientific, USA) according to manufacturer’s instructions.

### Phosphoproteomics

Scramble and siPKD1/2/3 transfected 3T3-L1 differentiated adipocytes treated with or without PMA (n=4 per group, 16 samples in total) were washed 3x with ice-cold trisaminomethane (Tris) buffered saline (TBS; 50 mM Tris-HCl and 150 mM sodium chloride pH 7.5 in Milli-Q water) and all residual TBS was aspirated after the final wash. Lysis was performed by adding 200 µl of ice-cold guanidinium chloride (GdmCl) lysis buffer (6 M GdmCl and 0.1 M Tris-HCl pH 8.5 in Milli-Q water) per well. Cells were then scraped, collected, and immediately boiled at 95°C for 5 min. Cells were then placed on ice to cool before sonication and lipid removal as above. An aliquot of protein was diluted 1:5 in 8 M urea and protein concentration was quantified via BCA assay. Samples were then diluted to equal protein concentrations in GdmCl lysis buffer.

Protein was precipitated with chloroform:methanol (1.25 volumes of 1:4), and resolubilised in sodium dodecyl sulfate (SDC) buffer (4% SDC/100 mM Tris pH 8.5) with sonication, and 250 µg was digested and processed through the EasyPhos workflow (30). Enriched phosphopeptides in MS loading buffer were analysed using an Orbitrap Exploris 480 mass spectrometer (Thermo Scientific, USA) interfaced to a Dionex U3000 HPLC with an in-house assembled 50-cm nanospray column with an inner diameter of 75 µm and packed with 1.9 µm C18 ReproSil Pur AQ particles (Dr. Maisch GmbH). Column temperature was maintained at 60°C using a column oven (Sonation, Germany), and peptides were separated with a binary buffer system comprising 0.1% formic acid (buffer A) and 80% acetonitrile in 0.1% formic acid (buffer B), at a flow rate of 400 nL/min, with a gradient of 3-19% buffer B over 40 min followed by 19-41% buffer B over 20 min, resulting in ∼1-hr gradients. Peptides were analysed in the mass spectrometer with a survey scan (350-1400 m/z, R = 120,000) at a target of 3e6 ions, followed by 48 data-independent acquisition (DIA) MS/MS scans (350-1022 m/z) with higher-energy collisional dissociation (HCD) (normalised collision energy 25%), at a target of 3e6 ions (max injection time 22 ms, isolation window 14 m/z, 1 m/z window overlap), and fragment ions were detected in the Orbitrap (R = 15,000). Raw data was processed using Spectronaut (v16.0.220606.53000). Data were searched using the directDIA analysis workflow, against a Mus musculus UniProt Reference Proteome database (March 2022 release). Key Spectronaut settings were: precursor and protein Qvalue cutoffs 0.01; Qvalue filtering; MS2-level quantification; fixed modifications “Carbamidomethyl (C)”, variable modifications “Acetyl (Protein N-term)”, “Oxidation (M)”, “Phospho (STY)”; imputation strategy “none”; cross run normalization “on” (automatic); PTM localization “True”; PTM Probability cut-off 0.5 (further filtering for Class 1 phosphosites with localization probability > 0.75 was performed during downstream analysis); multiplicity “on”; PTM consolidation “linear model”.

Statistical analysis was performed using the Perseus platform (v2.0.7.0) (31). Data was initially log_2_(x) transformed and the level of correlation of phosphosite enrichment between replicates across all groups was then calculated using Pearson’s correlation coefficient (r). Multiple comparison testing was then completed using a two-way analysis of variance (ANOVA; siRNA treatment, PMA treatment) with permutation-based false discovery rate (FDR) set to 0.05. Enriched phosphosites were filtered for ANOVA significance and z-scores were calculated for each biological replicate across significant phosphosites. A heatmap with Euclidean distance-based hierarchical clustering of enriched ANOVA-significant phosphosite z-scores was then generated.

Given that PMA treatment caused a dramatic change in the 3T3-L1 phosphoproteome, and that PKD isoform phosphosites were undetected in DMSO treated cells, the siPKD1/2/3-PMA and scramble-PMA groups were isolated for further analysis. Data was then filtered for phosphosites quantified in at least three biological replicates in at least one group. Two-sample Student’s t-tests were then performed for each phosphosite between siPKD1/2/3-PMA and scramble-PMA groups, with multiple comparison correction via Benjamini-Hochberg FDR set to 0.05. Substrate prediction was performed using PhosphoSitePlus (v6.7.4), with a log2 (score) of > 1.5 and site percentile > 90% considered a predicted site. For bulk pathway analysis, phosphosites with p<0.05 without multiple comparison testing were used. The list of significant sites was filtered for unique proteins, and PKD phosphosites were removed from analysis. The filtered list was then analysed in the Gene Ontology Protein Analysis Through Evolutionary Relationships (PANTHER) classification system (v17.0) (32) for overrepresented pathways in the Panther, Reactome and Gene Ontology (GO) biological process databases using Fisher’s exact test with FDR multiple comparison testing. The complete list of all phosphosites detected in 3T3L1 cells in this analysis was used as the reference list. No pathways were found to be statistically significantly overrepresented after FDR correction. Therefore, the top ten overrepresented pathways by unadjusted p-value were displayed, ranked by fold change. Dot plots for pathway analysis were generated using the R statistical analysis software (v4.4.1) with ‘ggplot2’ package (v3.5.1).

### Metabolic flux, glucose uptake and lipolysis

Metabolic flux assessment using the Seahorse XFe analyser and 2-[1,2-^3^H(N)]-deoxy-D-glucose (^3^H-2DOG; Perkin Elmer, USA) uptake assays were performed on scramble and siPKD1/2/3 transfected 3T3-L1 differentiated adipocytes five days post-differentiation in 24-well plates, as previously described (33). Lipolysis was assessed on scramble and siPKD1/2/3 transfected 3T3-L1 differentiated adipocytes 5 days post-differentiation in 24-well plates. Cells were washed 2x in Krebs–Ringer Phosphate (KRP) buffer (0.6 mM Na_2_HPO_4_, 0.4 mM NaH_2_PO_4_, 120 mM NaCl, 6 mM KCl, 1 mM CaCl_2_, 1.2 mM MgSO_4_, 12.5 mM HEPES, 5 mM glucose, pH 7.4) before being serum starved in DMEM supplemented with 0.02% fatty-acid free BSA for 2 hrs. Cells were then incubated for 1 hr in KRP supplemented with 3.5% fatty-acid free BSA, and either 10 nM insulin, and/or 1 nM CL316,243 (Sigma-Aldrich, USA), a selective β3-adrenergic receptor agonist that promotes adipose tissue lipolysis (34). After 1 hr, media and protein were collected and stored at –80°C. Media glycerol was determined as a measure of lipolysis using a glycerol assay kit (Sigma-Aldrich, USA) according to manufacturer’s instructions. Glycerol release was normalised to protein content of lysed cells, determined by BCA assay.

### Animals

All breeding and experimental procedures involving animals were approved by the Deakin University Animal Welfare Committee (G14-2018 and G15-2018), which is subject to the Australian Code for the Responsible Conduct of Research. Mice were group housed with 4-5 mice per cage with a 12-hr light/dark cycle at 22°C with ad libitum access to food and water. Mice were maintained on control chow diet (20% protein, 5% fat; Barastoc Rat and Mouse, Ridley Agriproducts, Australia) for the duration of the study.

A novel dominant negative (DN) PKD knock-in mouse model was generated as previously described (35). The DNPKD transgene was inserted at the *Rosa26* locus and was regulated by a floxed STOP codon flanked by loxP sites that prevents the expression of the DNPKD transgene. DNPKD mice were crossed with AdipoQ-CreERT2 mice, which express a tamoxifen activatable Cre-recombinase driven by the adipocyte-specific adiponectin promoter (36). To generate adipose specific DNPKD experimental mice, heterozygous DNPKD^-/+^ mice were paired with AdipoQ-CreERT2^+^ mice. This generated male and female AdipoQ-CreERT2^+^.DNPKD^-/-^ offspring, which were used as control mice. Heterozygous AdipoQ-CreERT2^+^.DNPKD^-/+^ mice were paired to generate homozygous AdipoQ-CreERT2^+^.DNPKD^+/+^ mice (atDNPKD). Control and atDNPKD mice were separated as littermates by no more than two generations. Experimental animals were age matched and were analysed as two distinct cohorts of roughly equal size. To induce DNPKD expression, tamoxifen (Sigma-Aldrich, USA) powder was resuspended in 100% ethanol to a concentration of 250 mg/ml and sonicated in a water bath at room temperature (RT) for 10 min. Tamoxifen solution was then diluted to a concentration of 25 mg/ml in soybean oil (Sigma-Aldrich, USA) pre-warmed to 37°C, then stored at 4°C. Tamoxifen-soybean oil solution was administered at RT once daily for 5 days at 100mg/kg to both male and female control and atDNPKD mice between 12 and 16 weeks of age. Mice underwent a 2-week wash-out period following tamoxifen administration prior to the start of experiments described below.

### Animal experiments

Body weight was recorded at baseline (2 weeks post-tamoxifen administration) and twice per week for the duration of the study. Body composition was assessed at baseline, prior to major experiments, and at the conclusion of the study by magnetic resonance imaging (EchoMRI™ Whole Body Composition Analyzer, EchoMRI, Singapore). Energy expenditure and substrate utilisation were measured via indirect calorimetry using Promethion™ metabolic analysis cages (Sable Systems, USA) linked to Promethion Live software (v21.0.3; Sable Systems, USA) in response to three experimental challenges as described in detail below. Raw data was processed in ExpeData analysis software (v1.9.27; Sable Systems, USA) using in-built macros (‘Macro 1’ followed by ‘Macro 13’). Total activity was calculated by the addition of all x, y, and z plane laser break counts. VO_2_ and VCO_2_ data was then used to calculate rates of energy expenditure, glucose oxidation, and lipid oxidation using previously validated formulas as shown below (37, 38).

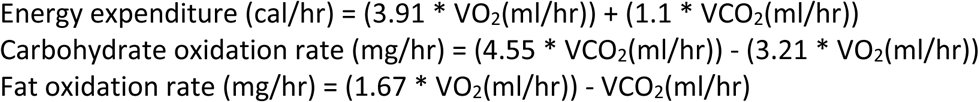

For basal energy expenditure, control and atDNPKD mice at 7 weeks post-tamoxifen administration were placed in metabolic cages and data were recorded for 25 hrs. Data from the first hour was disregarded as mice adjusted to the new environment. Data was calculated and analysed as hourly averages for 24 hrs. To characterise responses to a β-adrenergic stimulation, control and atDNPKD mice at 8 weeks post-tamoxifen administration were placed in metabolic cages for 2 hrs. Mice were then administered the specific β_3_-adrenergic receptor agonist, CL316,243 (Sigma-Aldrich, USA) at 1 mg/kg lean mass in sterile Milli-Q water via intraperitoneal injection and data were recorded for 4 hrs. Data from 30 min prior to CL316,243 administration and the final 30 min after CL316,243 administration were quantified. To assess substrate utilisation and food intake during fasting and refeeding, control and atDNPKD mice at 10 weeks post-tamoxifen administration were fasted overnight for 17 hr. Data in the final 4 hr of the fast and the first 4 hr of the refeed period were analysed. Analysis of covariance (ANCOVA) was performed on indirect calorimetry data with body mass as a covariate, as previously described (39), using the NIDDK Mouse Metabolic Phenotyping Center Energy Expenditure Analysis page (http://www.mmpc.org/shared/regression.aspx; supported by grants DK076169 and DK115255) (40–42).

An insulin tolerance test (ITT; 0.75 IU/kg lean mass – Actrapid; Novo Nordisk, Denmark) was performed at 12 weeks post-tamoxifen administration following a 5 hr fast. Blood glucose was measured via the tail vein using the Accu-Chek Performa Meter (Roche, Australia) at baseline and 20, 40, 60, 90 and 120 min post insulin administration. Blood (30 uL) was also collected via the tail vein at baseline, 20 and 40 min in heparinised tubes and centrifuged at 10 000 g for 2 min to separate plasma. Plasma was then stored at –20°C for later analysis of free fatty acid (FFA) concentration as a measure of lipolysis.

At 13 weeks post-tamoxifen administration, mice were fasted for 5 hrs and baseline fasting blood glucose was measured and blood (30 uL) was collected via the tail vein and placed in heparinised tubes. Mice were then administered CL316,243 at 1 mg/kg lean mass via intraperitoneal injection (43) to examine β-adrenergic stimulated lipolysis. Mice were humanely killed 15 min post-CL316,243 administration, and blood was immediately collected via cardiac puncture and placed in heparinised tubes. Blood samples from baseline and 15 min post-CL316,243 administration were centrifuged at 10 000 g for 2 min to separate plasma. Plasma was then stored at –20°C for later analysis of FFA concentration as a measure of lipolysis, and fasting insulin concentration to determine Homeostatic Model Assessment for Insulin Resistance (HOMA-IR).

### Plasma biochemistry

Plasma insulin was quantified via ELISA in plasma samples collected during the ITT and at experimental endpoint as previously described (44). Fasting insulin in pmol/L was then converted to µIU/mL using a conversion factor of 1 µIU/mL = 6 pmol/l (45). HOMA-IR was then calculated using the following equation (46):

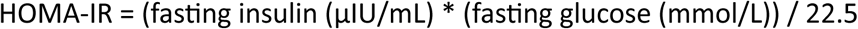

Plasma FFA concentration was determined using the NEFA-C kit (Wako, Japan) with a modified protocol for use in 96-well plates. Assay standard was diluted in Milli-Q water to concentrations of 1, 0.8, 0.5, 0.25, 0.1 and 0.05 mM to generate a standard curve. Standards, water blank, and plasma samples (5 µL) were loaded into a clear 96-well plate. Reagent 1 ( 65 µL) was added, and the plate was incubated at 37°C for 10 min. Reagent 2 (130 µL) was then added, and the plate was incubated at 37°C for 15 min. Absorbance was then read on an xMark™ microplate spectrophotometer (Bio-Rad, USA) at 550 nm and FFA concentration was calculated using the standard curve.

### Gene and protein expression

For gene expression analysis, scramble and siPKD1/2/3 transfected 3T3-L1 differentiated adipocytes were washed in ice-cold PBS and lysed in RNeasy lysis buffer RLT (Qiagen, Germany) containing 1% β-mercaptoethanol (Sigma-Aldrich, USA). An equal volume of 70% ethanol was then added, and RNA was extracted using the RNeasy Mini Kit (Qiagen, Germany) according to manufacturer’s instructions. RNA was reverse transcribed to generate cDNA using the Maxima H Minus First Strand cDNA synthesis kit (Thermo Scientific, USA) as per manufacturer’s instructions in a SimpliAmp thermal cycler (Thermo Scientific, USA). cDNA was then diluted 1:10 in nuclease free water (Thermo Scientific, USA). Diluted cDNA was quantified using the Quant-iT OliGreen ssDNA Assay Kit (Thermo Scientific, USA) as per manufacturer’s instructions in a Flexstation II^384^ microplate reader (Molecular Devices, USA) set to 10 s plate shake, 485 nm excitation and 538 nm emission.

Gene expression was quantified in 1 µL of cDNA via real-time quantitative polymerase chain reaction (RT-qPCR) using the Luminaris HiGreen qPCR master mix (Thermo Scientific, USA) and specific forward and reverse primers for target genes (Table 1) on a PikoReal 96 RT-qPCR system (Thermo Scientific, USA). The qPCR protocol included 10 min at 95°C, then 40 cycles of 95°C for 30 s and 60°C for 60 s. Relative mRNA expression was then calculated using the cycle threshold (C_T_) relative quantification method (47) with cyclophilin gene expression used as a housekeeping gene.

**TABLE 1:**
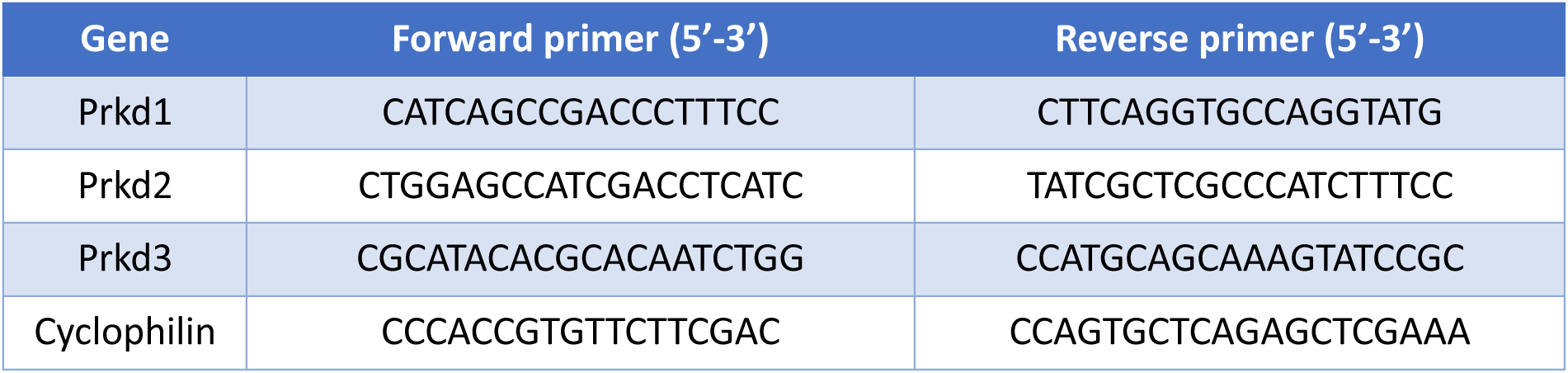
Target genes and associated primers used in real-time quantitative PCR.

Protein expression was measured via immunoblotting. Lysates from scramble and siPKD1/2/3 transfected 3T3-L1 differentiated adipocytes were diluted to equal protein concentrations according to BCA assay in protein lysis buffer before the addition of solubilizing buffer (0.125 M Tris-HCl pH 6.8, 4% SDS, 10% glycerol, 4 M urea, 10% β-mercaptoethanol, and 0.001% bromophenol blue). For adipose tissue analysis, 20-40 mg of tissue was homogenised in ice cold protein lysis buffer using a handheld homogeniser, before undergoing the sonication and lysate clearance procedure outlined previously for 3T3-L1 adipocytes. Protein (5-10 µg) was separated by sodium dodecyl sulphate polyacrylamide gel electrophoresis (SDS-PAGE) (Bio-Rad, USA) in 26-well 4–15% Criterion™ TGX Stain-Free™ Protein Gels (Bio-Rad, USA) at 200 V for 60 min. Protein was transferred onto nitrocellulose membrane (Bio-Rad, USA), which was incubated in Miser™ Antibody Extender Solution NC (Thermo Scientific, USA) for 10 min. The membrane was then incubated overnight at 4°C in primary antibody diluted in TBST targeted against αTubulin (Sigma-Aldrich; T6074), CREB (Abcam; ab32515), CREB pSer^133^ (Cell Signaling; 9198), PKD1 (Abcam; ab108963), PKD3 (Cell Signaling; 5655), PKD pSer^744/748^ (Cell Signaling; 2054), and PKD pSer^916^ (Cell Signaling; 2051). Blots were visualised using the Clarity™ Western ECL Blotting Substrates kit (Bio-Rad, USA) or SuperSignal™ West Femto Maximum Sensitivity Substrate (Thermo Scientific, USA) in the ChemiDoc™ XRS+ and quantified with Image Lab software (v6.0.1 standard edition; Bio-Rad, USA). Protein abundance was quantified relative to αTubulin as a loading control, and expressed as the fold-change relative to control.

### Statistics

All data, except phosphoproteomic experiments as described above, was analysed using GraphPad Prism using independent samples t-tests or two-way ANOVA unless otherwise indicated. Post-hoc analysis was performed using Tukey’s multiple comparison test when a significant interaction effect was found from ANOVA analysis. Statistical significance was set at p<0.05. Data presented as mean ± SEM with individual data points represented, unless otherwise specified.

## RESULTS

### Identification of PKD-regulated signalling networks in 3T3-L1 adipocytes

To identify PKD regulated signalling networks in adipocytes, a PKD loss-of-function model was developed using a siRNA-mediated knockdown approach in 3T3-L1 adipocytes (Supplementary Figure 1A). The redundancy between PKD isoforms in this model was assessed by examining the phosphorylation status of both PKD and the downstream substrate CREB at serine 133 (Ser^133^) following exposure to PMA, a non-specific PKD activator. Simultaneous knockdown of all three PKD isoforms was required to abrogate PKD signalling (Figure 1A). This model was used to identify potential PKD-regulated phosphorylation by phosphoproteomic analyses using the EasyPhos platform (30). Initial validation of these samples by western blotting showed that simultaneous PKD1/2/3 knockdown reduced the phosphorylation of PKD regulatory sites at Ser^916^ and Ser^744/748^ (Supplementary Figure 1B). The reproducibility of the EasyPhos workflow was measured between biological replicates using Pearson’s correlation analysis. There was an average Pearson’s correlation coefficient (r) of 0.93 between replicates in the same group, and 0.90 in replicates between conditions (Supplementary Figure 1C). More than 16,500 class 1 phosphosites were identified across all samples and groups. Using ANOVA analysis with a permutation-based false discovery rate (FDR) set to p<0.05, 3,374 phosphosites (∼20% of the total phosphoproteome) were found to be differentially regulated across at least one of the four conditions. Hierarchical clustering of z-scores revealed that the majority of identified phosphosites were regulated by PMA treatment (Figure 1B).

**Figure 1:**
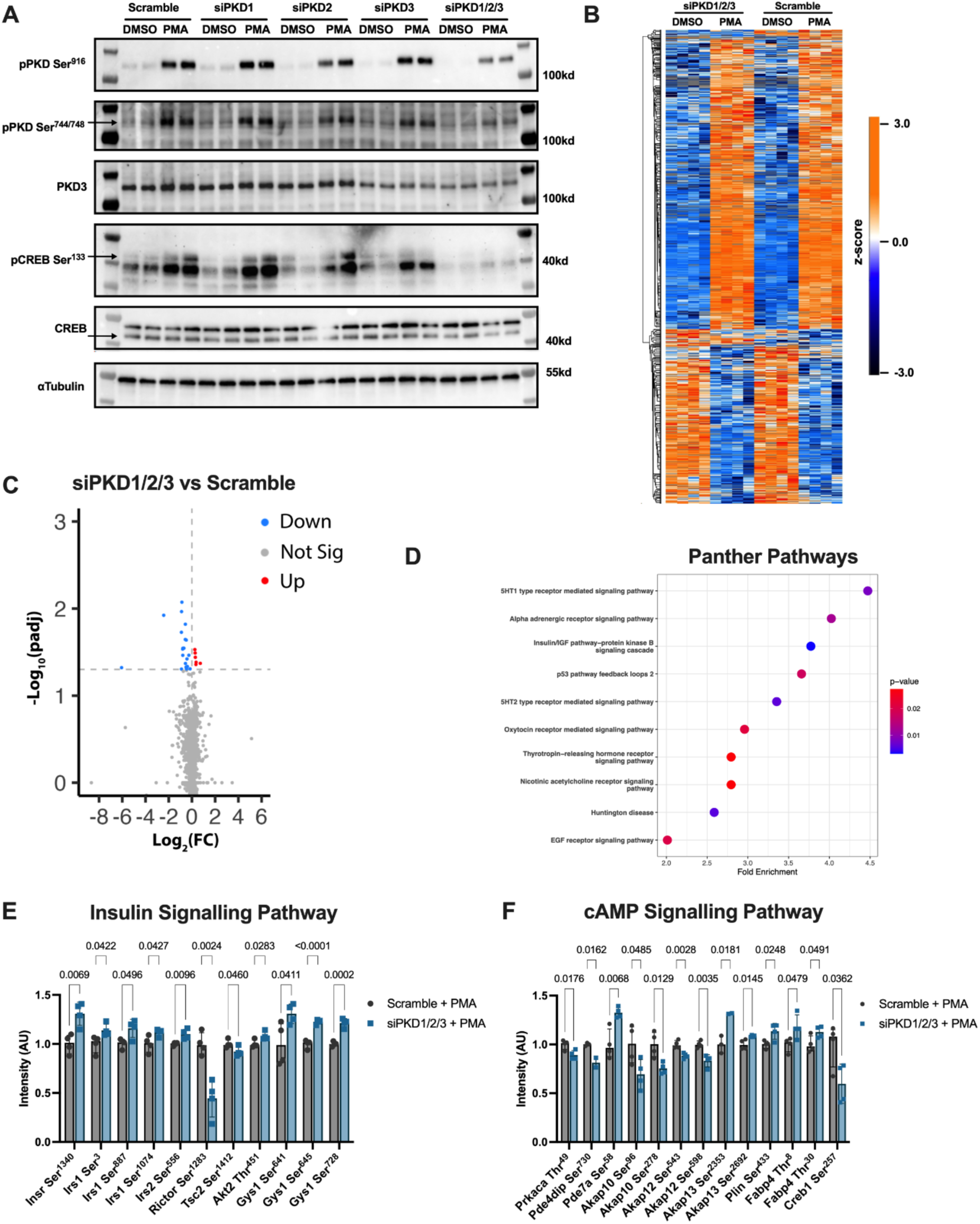
Phosphoproteomics analysis reveals metabolic regulation downstream of protein kinase D activation in differentiated 3T3-L1 adipocytes. Protein kinase D (PKD) and downstream cAMP response element-binding protein (CREB) protein expression via western blot with αTubulin as loading control in scramble and siPKD1/2/3 adipocytes; n = 2/group (A). Euclidean distance-based hierarchical clustering of enriched ANOVA-significant phosphosite z-scores (B). Volcano plot displaying the –Log_10_ false discovery rate (FDR) adjusted p-value and the Log_2_ fold change in intensity of phosphosites between PMA treated scramble and siPKD1/2/3 adipocytes (C). Top ten overrepresented Panther pathways by unadjusted p-value significance ranked by fold change identified in the 432 differentially regulated phosphosites before multiple comparison testing (D). Extracted phosphosites involved in insulin (E) and cAMP (F) signalling pathways displayed as mean ± SEM signal intensity. n = 4/group for phosphoproteomics experiments. Displayed p-values obtained by independent samples t-test.

To first validate sufficient PKD knockdown, individual PKD phosphosites were analysed between scramble and PKD1/2/3 knockdown cells following PMA treatment. The enrichment of PKD2 Ser^711^ and PKD3 Ser^734^, Ser^216^, Ser^213^ and Ser^30^ phosphopeptides were all significantly reduced by PKD1/2/3 knockdown (Supplementary Figure 1D). There were trends for reduced PKD2 Ser^873^ and PKD3 Ser^27^ (p = 0.056 and 0.081, respectively; Supplementary Figure 1D). PKD1 phosphopeptides were not detected in these samples. PKD-phosphopeptides were completely absent across all vehicle samples and was only present in PMA-treated samples. As such further analysis was performed between the PMA-treated control and PKD1/2/3 knockdown groups. This identified 432 differentially regulated phosphosites in PKD1/2/3 samples versus control before multiple comparison testing (Figure 1C). There were 26 phosphosites that remained significantly different after multiple comparison testing, with the majority downregulated by PKD1/2/3 knockdown following PMA stimulation (Figure 1C and Table 2). Some regulated phosphosites were predicted to be direct substrates of PKD, while others that were not predicted PKD substrates are likely regulated through secondary mechanisms (Table 2). Many differentially regulated phosphopeptides were from proteins involved in intracellular vesicular transport and cytoskeletal rearrangement, including Tbc1d22b, Itsn1, Tbc1d5, Dock7 and Stard13, which are verified biological functions regulated by PKD (7, 12, 19). However, the majority of identified downstream proteins had not previously been implicated in PKD signalling.

**TABLE 2:**
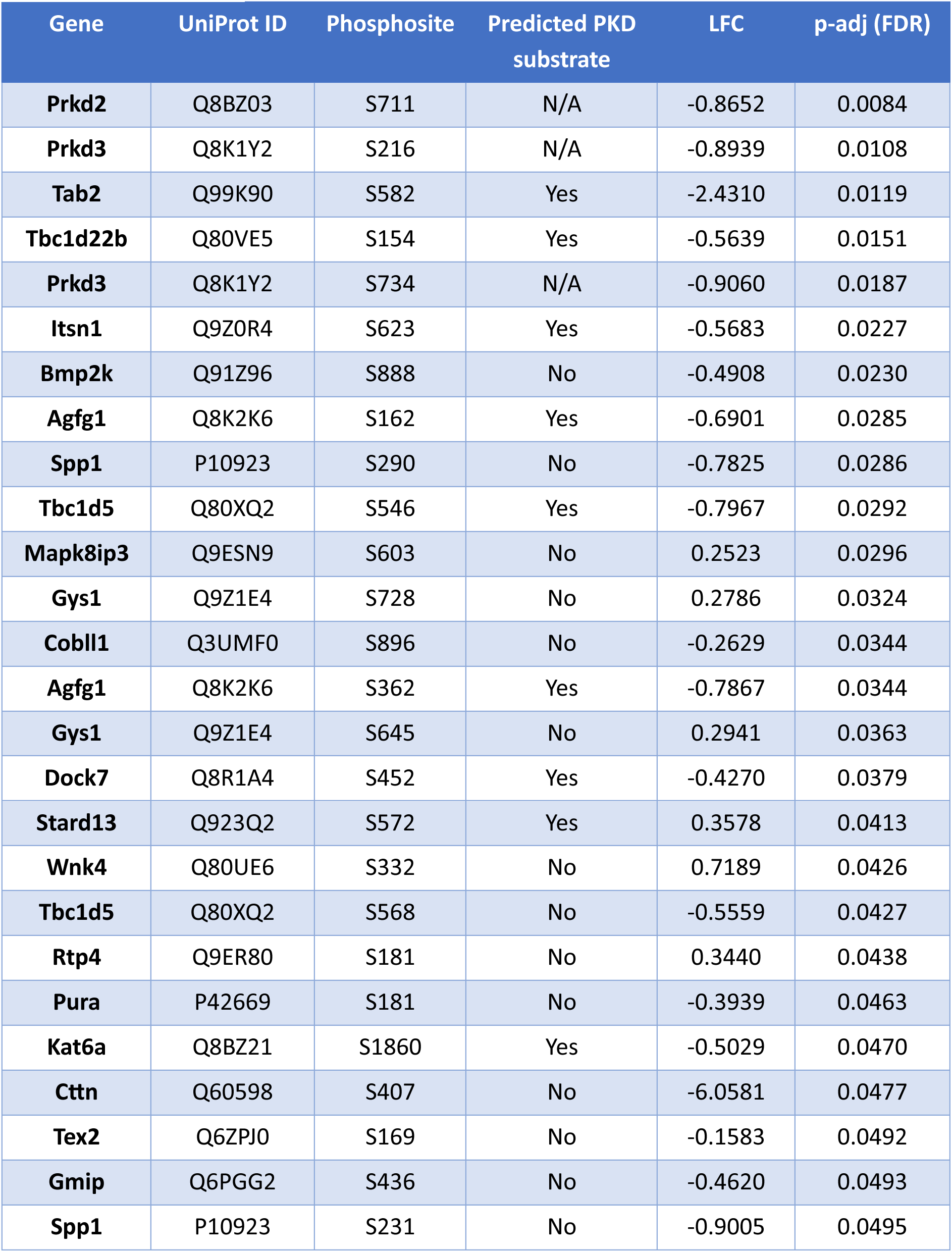
Downstream targets of protein kinase D in differentiated 3T3-L1 adipocytes. Differentially regulated phosphosites in response to PKD1/2/3 knockdown in PMA treated adipocytes ranked by statistical significance. LFC = the log_2_ fold change expression when compared to scramble siRNA treated adipocytes. P-value obtained by independent samples t-test adjusted for multiple comparison testing via false discovery rate (FDR). Substrate prediction was performed using PhosphoSitePlus. N/A denotes not applicable. n = 4 biological replicates/group.

To gain further insight into the molecular functions of these PKD-regulated phosphosites, pathway analysis was performed using the 432 phosphosites differentially regulated before multiple comparison testing. This again revealed enrichment of pathways normally associated with PKD activation, such as regulation of the actin cytoskeleton, tight junctions, and proteoglycan regulation (Figure 1D and Supplementary Figure 2B-C). Additional enriched pathways were those involved in metabolic regulation, including pathways of endocrine action, including serotonin, insulin, growth hormone and adrenergic signalling, as well as pathways related to metabolic regulation (Figure 1D and Supplementary Figure 2B-C).

Many of these pathways share common targets from the insulin signalling and the cAMP signalling pathways, which are integral to metabolic processes in adipocytes. Indeed, two of the top three overrepresented Panther pathways by fold change were ‘Insulin/IGF pathway-protein kinase B signalling cascade’ and ‘Alpha adrenergic receptor signalling pathway’ (Figure 1D). Therefore, the 432 phosphosites differentially regulated before multiple comparison testing were next interrogated for components of the insulin signalling pathway (Figure 1E). Under PMA stimulated conditions, knockdown of PKD1/2/3 generally altered phosphosites in a way that suggested inhibition of this pathway. For example, knockdown of PKD1/2/3 increased the phosphorylation of serine sites on IRS1 and 2 that are associated with Akt inactivation (48, 49). Similarly, knockdown of PKD1/2/3 increased the phosphorylation of inhibitory sites on glycogen synthase that are typically phosphorylated by GSK3b (50), despite adipocytes storing little to no glycogen.

Interrogation of the 432 differentially regulated phosphosites for components of the cAMP pathway, which regulates lipolysis and energy expenditure in adipocytes, identified phosphosites in proteins including the catalytic subunit of PKA, phosphodiesterases, Akap proteins, and proteins involved in lipolytic processes, such as perilipin and FABP4 (Figure 1F). The functional role of many of these phosphosites is unknown and most were not predicted PKD target sites based on sequence motif prediction. In both the insulin and cAMP pathways, the effect of PKD1/2/3 knockdown on these phosphosites with PMA stimulation was relatively subtle, possibly indicating that the role of PKD is to fine tune biological processes in adipocytes. Nonetheless, these data suggest that PKD influences key signalling networks in adipocytes related to metabolic regulation.

### Metabolic effects of PKD1/2/3 knockdown in 3T3-L1 adipocytes

As the insulin and cAMP signalling pathways reciprocally regulate adipocyte lipolysis, the impact of PKD1/2/3 knockdown on lipolysis was examined in 3T3-L1 adipocytes. Insulin and CL316,243, a specific β_3_ adrenergic receptor agonist that increases cAMP, suppressed and activated lipolysis as measured by glycerol release, respectively. However, there was no effect of PKD1/2/3 knockdown on either measure (Figure 2A). The effects of PKD1/2/3 knockdown on insulin-stimulated glucose uptake were also assessed, as PKD is known to regulate glucose uptake in some cell types through vesicular transport of plasma membrane glucose transporters (35, 51, 52), and insulin related pathways were identified by phosphoproteomics. However, there were no effects of PKD1/2/3 knockdown on either basal, insulin-stimulated and/or PMA-stimulated glucose uptake (Figure 2B).

**Figure 2:**
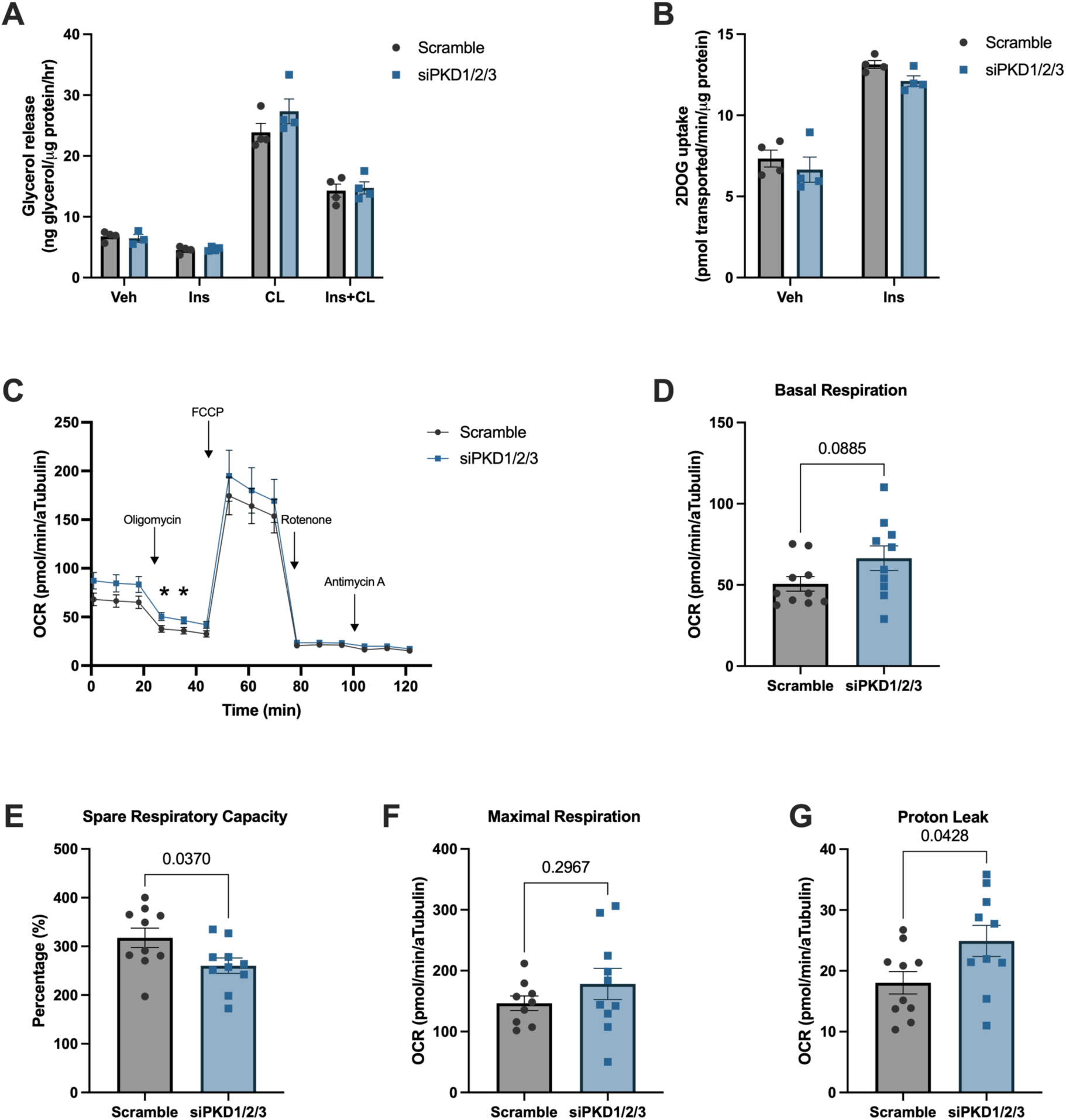
Loss of PKD isoforms increases basal energy expenditure and uncoupled respiration in differentiated 3T3-L1 adipocytes. Glycerol release **(A)** during a lipolysis assay and 2-deoxy-D-glucose uptake **(B)** during a glucose uptake assay. n = 4/group. Oxygen consumption rate (OCR) during a mitochondrial stress test **(C)** with calculated basal respiration **(D)**, Spare respiratory capacity as a percentage of basal respiration **(E)**, maximal respiration **(F)**, and proton leak **(G)**. OCR was corrected for αTubulin protein expression measured in cell lysates via western blot at the conclusion of the experiment. n = 10/group. Veh = vehicle; Ins = insulin; CL = CL316,243. Data are mean ± SEM. Displayed p-values obtained by independent samples t-test. * indicates p < 0.05 compared to scramble.

Activation of cAMP signalling in 3T3-L1 adipocytes also increases uncoupled respiration (53), which occurs in beige and brown adipose tissue *in vivo* leading to increased energy expenditure (54). Although adipocyte respiratory responses were insensitive to CL316,243 (data not shown), knockdown of PKD1/2/3 had subtle effects on oxidative energetics (Figure 2C). There was a trend for increased basal respiration (Figure 2D), and lower spare respiratory capacity as a percentage of basal respiration (Figure 2E). Given no differences in maximal respiration between groups (Figure 2F), a higher spare respiratory capacity percentage indicates a higher basal energy expenditure in siPKD1/2/3 adipocytes (55). This increase in respiration is likely due, at least in part, to an increase in uncoupled respiration, as observed by the increased proton leak in siPKD1/2/3 cells compared to control (Figure 2G). Overall, this evidence suggests that knockdown of PKD1/2/3 increases energy expenditure in adipocytes via uncoupled respiration but does not influence single end-point measures of lipolysis or glucose uptake downstream of insulin or cAMP pathway stimulation in 3T3-L1 adipocytes.

### Energy balance in adipose tissue dominant negative PKD mice

Following initial findings that PKD regulates key metabolic signalling pathways and promotes energy expenditure in adipocytes *in vitro*, the role of PKD in adipose tissue was further explored *in vivo* using atDNPKD mice. These mice overexpress a kinase dead PKD mutant isoform, specifically in adipose tissue, that outcompetes endogenous PKD to functionally inhibit PKD activity, with activity inhibition directly proportional to the extent of transgene overexpression (35). DNPKD induction can be visualised via western blot as a separate, higher molecular weight band using a PKD antibody (Figure 3A), and semi-quantified by an increase in total PKD protein expression. Successful induction of the DNPKD transgene in male (Figure 3 top panel) and female (Figure 3 bottom panel) mice in both visceral (vWAT; Figure 3A and 3D) and subcutaneous (sWAT; Figure 3B and 3E) white adipose tissue, and brown (BAT; Figure 3C and 3F) adipose tissue depots was observed, although total PKD protein expression was not significantly increased (p = 0.065) in sWAT of male mice (Figure 3B). To confirm adipose tissue specificity, PKD protein was analysed in liver, heart and skeletal muscle (Supplementary Figure 3). No DNPKD band or differences in total PKD protein expression were observed between control and atDNPKD mice, indicating that the induction of DNPKD expression was specific for adipose tissues.

**Figure 3:**
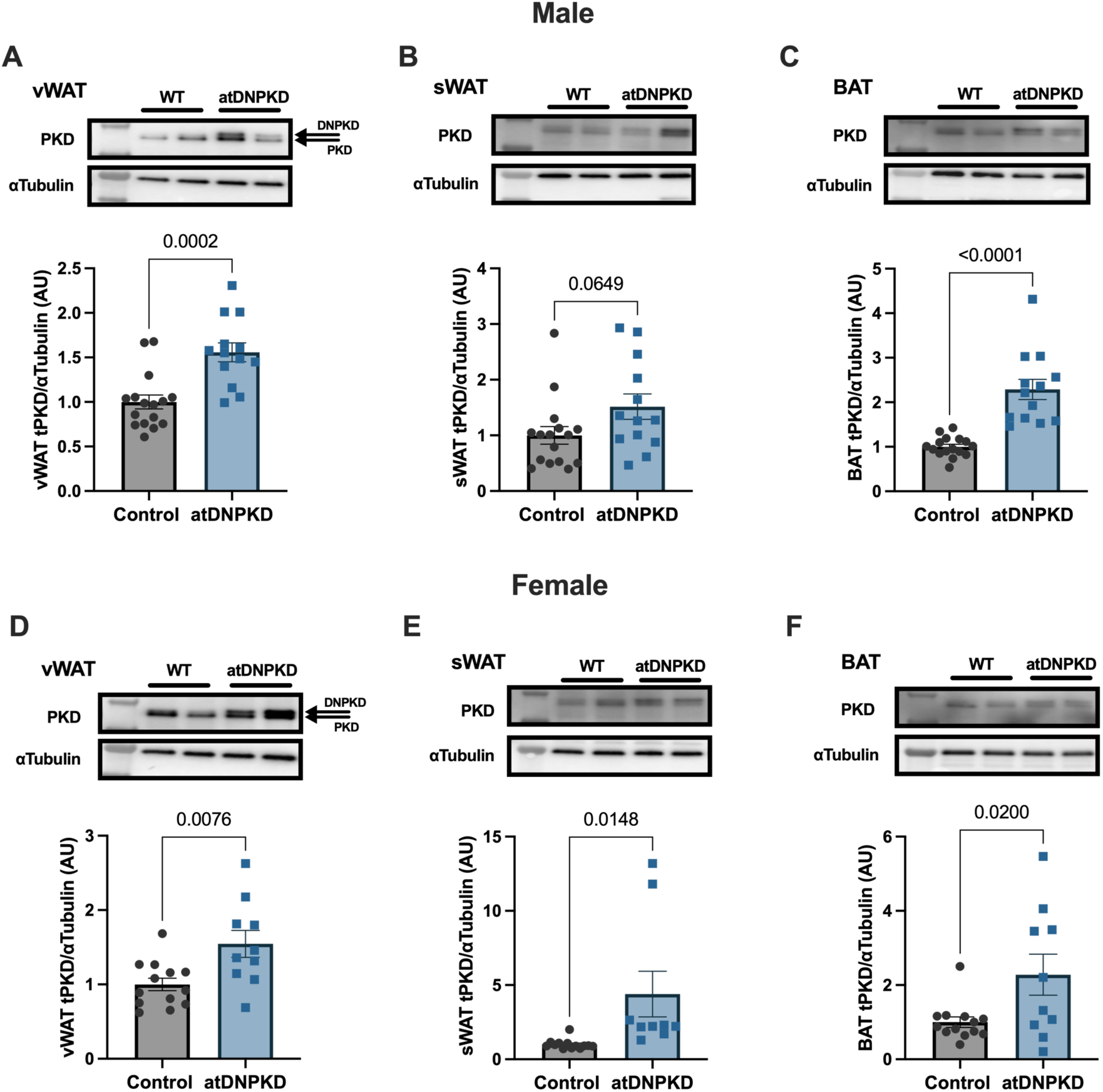
Adipose tissue dominant negative protein kinase D (DNPKD) transgene induction in male (A-C) and female (D-F) wild type (WT) and atDNPKD mice. Representative western blots and densitometric quantification of total protein kinase D (tPKD) expression in mouse visceral white adipose tissue (vWAT; **A and D**), subcutaneous white adipose tissue (sWAT; **B and E**), and brown adipose tissue (BAT; **C and F**). αTubulin used as protein loading control. AU = arbitrary unit. Data are mean ± SEM. n = 9-16/group. Displayed p-values obtained by independent samples t-test.

Body composition and basal energy balance was first assessed. In male mice, expression of DNPKD in adipose tissue had no effect on body mass (Figure 4A) but resulted in reduced lean mass at baseline and at the end of the experiment (Figure 4B). There were no differences in fat mass (Figure 4C). Furthermore, expression of DNPKD in adipose tissue had no effect on 24 hr energy expenditure (Figure 4D), food intake (Figure 4E) or substrate utilisation (Figures 4F and G). In female mice, expression of DNPKD in adipose tissue had no effect on body mass (Figure 4H) and although lean mass was higher in atDNPKD mice at baseline, there was no difference at the end of the experiment (Figure 4I). There were no differences in fat mass throughout the experiment (Figure 4J), nor any differences in energy expenditure, food intake or substrate utilisation (Figures 4K-N).

**Figure 4:**
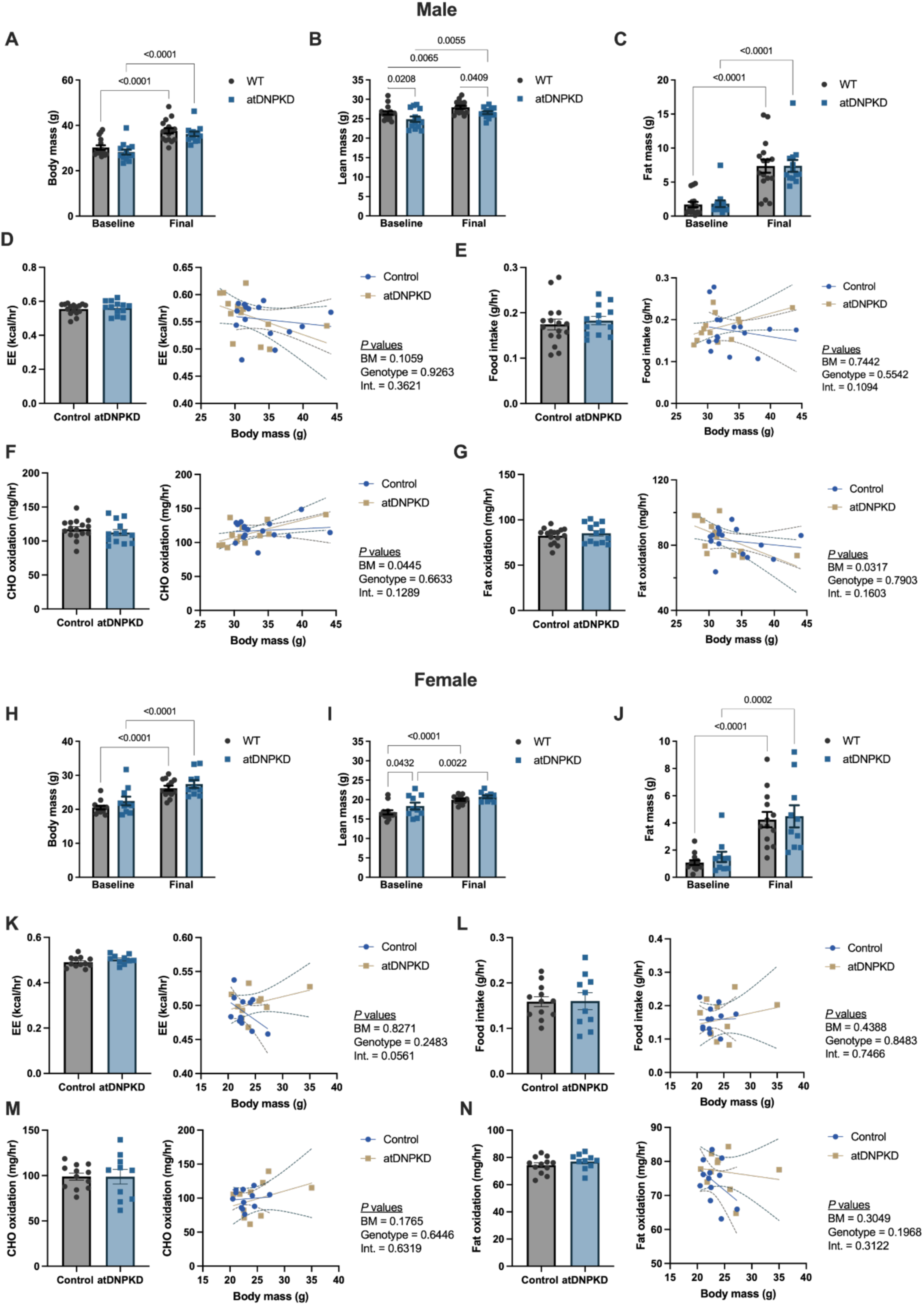
DNPKD expression in adipose tissue does not affect body composition and basal energy expenditure in male (A-G) and female (H-N) mice. Body mass **(A and H)**, lean mass **(B and I)**, and fat mass **(C and J)** were recorded two weeks-post tamoxifen administration (Baseline) and 13 weeks post tamoxifen administration (Final). At seven weeks post tamoxifen, mice were placed in metabolic cages and the average energy expenditure (EE; **D and K**), food intake **(E and L)**, carbohydrate (CHO) oxidation **(F and M)**, and fat oxidation **(G and N)** were quantified and analysed with body mass as a covariate via ANCOVA. Data are mean ± SEM. n = 10-16/group. Displayed p-values obtained by 2-way ANOVA with Fisher’s LSD post-hoc test or independent samples t-test.

### Insulin action in adipose tissue dominant negative PKD mice

As phosphoproteomic data in 3T3-L1 cells indicated a role for adipose PKD in the regulation of the insulin signalling pathway, the role of adipose tissue PKD on insulin-regulated glucose and fatty acid metabolism *in vivo* was investigated. In male mice, there were no differences in fasting blood glucose (Figure 5A), plasma insulin (Figure 5B) or insulin sensitivity, estimated by HOMA-IR (Figure 5C). Furthermore, there was no difference in blood glucose kinetics during an insulin tolerance test (Figure 5D). Fasting plasma FFA were elevated at baseline and remained increased 20 min after administration of insulin in atDNPKD mice, whereas plasma FFA was significantly reduced at 20 min post-insulin in control mice (Figure 5E). By 40 min after insulin administration, there were no differences between control and atDNPKD mice, however, the FFA area under the curve was increased in atDNPKD mice (Figure 5F). Despite opposing changes in plasma FFA in response to insulin at 20 min in control and atDNPKD mice, the change in FFA concentration 20 min after insulin administration was not statistically different between groups (Figure 5G).

**Figure 5:**
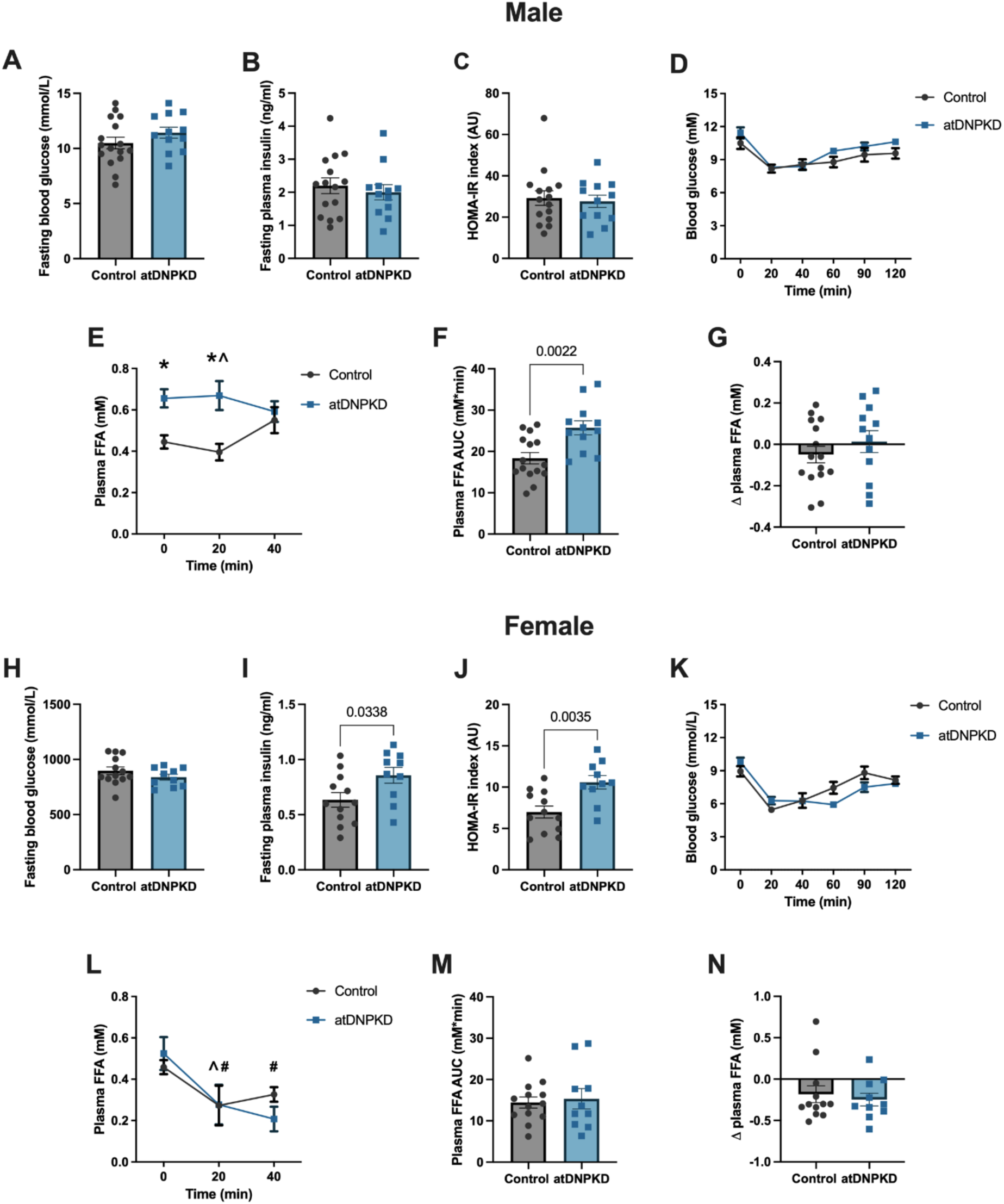
DNPKD expression in adipose tissue does not alter metabolic responses to insulin in male (A-G) and female (H-N) mice. Fasting blood glucose **(A and H)**, fasting plasma insulin **(B and I)** and HOMA-IR index **(C and J)**. Blood glucose **(D and K)**, plasma free fatty acids (FFA) **(E and L)** and plasma FFA area under the curve (AUC; **F and M**) during an insulin tolerance test (ITT; 0.75 IU/kg lean mass). Change in plasma FFA in the first 20 min of the ITT **(G and N)**. Data are mean ± SEM. n = 10-16/group. Displayed p-values obtained by independent samples t-test. *indicates p < 0.05 compared to Control. ^indicates p<0.05 vs 0 min in Control mice. #indicates p<0.05 vs 0 min in atDNPKD mice.

In female mice, there were no differences in fasting blood glucose between groups (Figure 5H), however female atDNPKD mice had significantly higher fasting plasma insulin (Figure 5I). This resulted in impaired insulin sensitivity predicted by HOMA-IR (Figure 5J). However, glucose kinetics were not different between control and atDNPKD mice following insulin administration (Figure 5K). There were also no differences in fasting plasma FFA, nor plasma FFA following insulin administration (Figures 5L and M). There were no differences in the suppression of plasma FFA between groups, 20 min after insulin administration (Figure 5N). These data suggest that adipose tissue PKD is important for fatty acid homeostasis in male mice. However, consistent with our *in vitro* data, there appears to be no role for adipose tissue PKD in regulating acute insulin action on glucose and fatty acid homeostasis.

### Metabolic responses to β_3_-adrenoceptor agonism in adipose tissue dominant negative PKD mice

Several pathways regulated by PKD identified through phosphoproteomics included cAMP signalling, which is typically engaged in adipose tissue during fasting or cold exposure through adrenergic stimulation (56). β-adrenergic signalling is the predominate adrenergic pathway in mouse adipose tissue (57). Therefore, the metabolic responses to the β_3_-adrenergic receptor agonist CL316,243 were examined. In male mice, plasma FFAs were higher in atDNPKD mice following CL316,243 administration (Figure 6A). Interestingly, the higher fasting plasma FFA concentration previously observed in male atDNPKD mice compared to controls (Figure 5D) was not seen in this experiment (Figure 6A). Nonetheless, the change in plasma FFAs following CL316,243 administration was greater in atDNPKD mice (Figure 6B), suggesting greater sensitivity to β3-adrenergic receptor agonism. This was further explored in other aspects of metabolic homeostasis. The increase in energy expenditure following CL316,243 administration tended to be greater (p=0.0642) in atDNPKD mice, which was statistically significant in an ANCOVA analysis with body mass as a covariant (Figure 6C). Interestingly, a significant interaction was also observed, suggesting this effect was most pronounced in mice with lower body mass (Figure 6C). There were no differences in carbohydrate oxidation between groups following CL316,243 administration (Figure 6D). However, the increase in lipid oxidation following CL316,243 administration tended to be higher in atDNPKD mice and ANCOVA analysis replicated the significant changes observed for energy expenditure (Figure 6E). These data suggest that adipose tissue PKD inhibits energy expenditure and fat oxidation in response to adrenergic stimulation in male mice, and this effect diminishes as body weight increases. These responses were also examined in female mice and there were no differences in plasma FFAs (Figure 6F and 6G), energy expenditure (Figure 6H), and substrate utilisation (Figures 6I and 6J) following CL316,243 administration. Taken together, these data suggest that adipose tissue PKD is an inhibitory checkpoint for adrenergic signalling in the regulation of lipolysis, fat utilisation and energy expenditure in male mice, but not in female mice.

**Figure 6:**
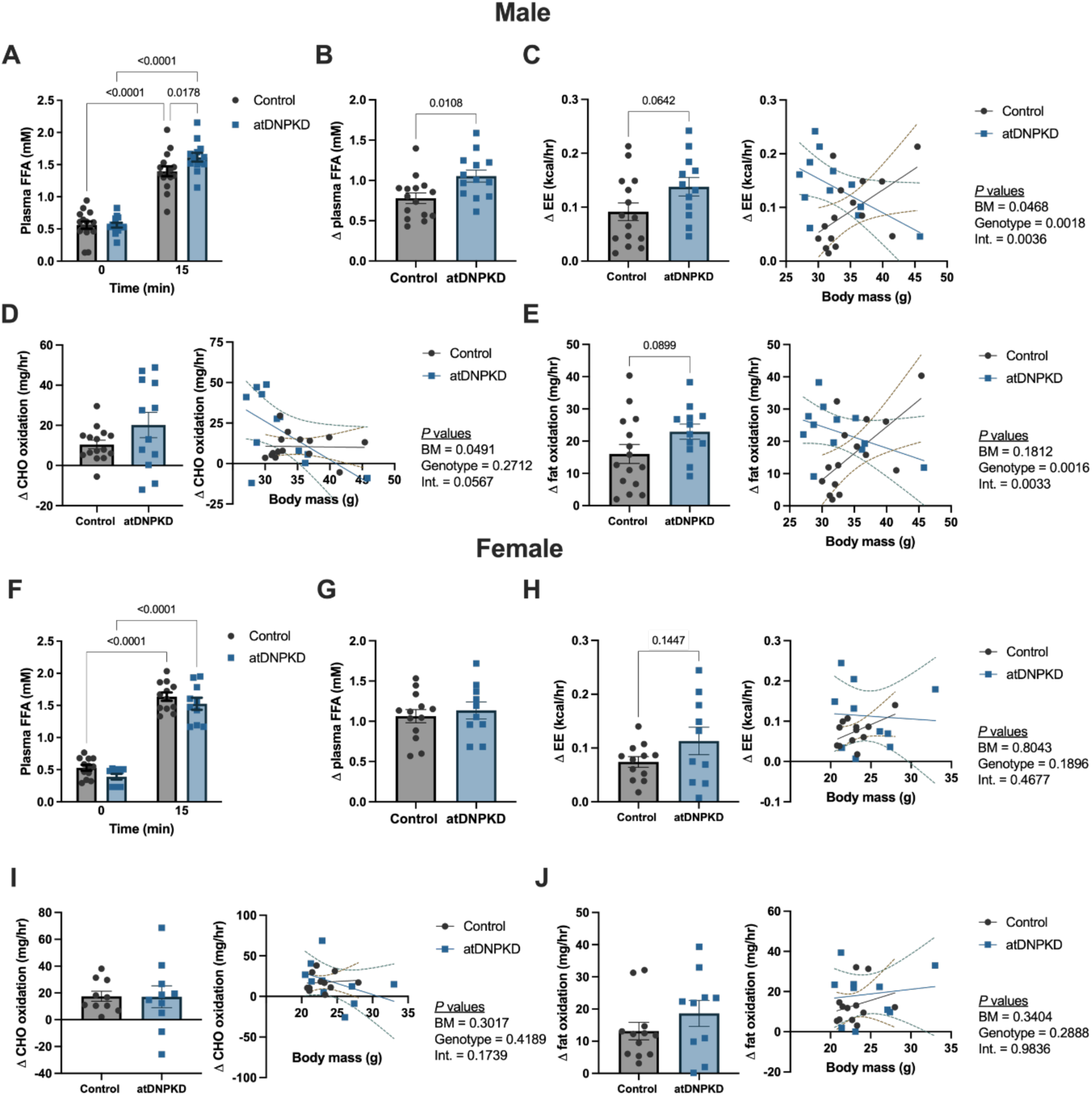
DNPKD expression in adipose tissue potentiates β-adrenergic signalling in male (A-E) but not in female (F-J) mice. Plasma free fatty acids (FFA) **(A and F)** and the change (Δ) in plasma FFA **(B and G)** before and 15 min after administration of the selective β-adrenergic agonist, CL316,243 (1 mg/kg lean mass). The Δ energy expenditure **(C and H),** carbohydrate (CHO) oxidation **(D and I)** and fat oxidation **(E and J)** before and 4 hrs after administration of CL316,243. Data are mean ± SEM. n = 10-16/group. Displayed p-values obtained by independent samples t-test.

### Energy expenditure and substrate utilisation during fasting in adipose tissue dominant negative PKD mice

The previous experiments suggest that adipose tissue PKD could be important for restraining fat utilisation and energy expenditure, potentially to maintain body weight, in response to metabolic challenges such as fasting. To test this hypothesis, atDNPKD mice were housed in metabolic cages during an overnight fast and metabolic responses in the final 4 hr of the fast were analysed. In male mice, there were no differences in fasting energy expenditure between groups (Figure 7A). Fat oxidation was higher in atDNPKD mice when assessed with body weight as a covariant in ANCOVA analysis (Figure 7B). There were no differences in fasting carbohydrate oxidation (Figure 7C), nor any difference in the change in body weight (Figure 7D) during fasting between groups. In female mice, there were no differences in fasting energy expenditure (Figure 7E), substrate utilisation (Figure 7F and 7G), and change in body weight (Figure 7H). These data indicate that adipose tissue PKD regulates fat oxidation during fasting in male mice, but not in female mice.

**Figure 7:**
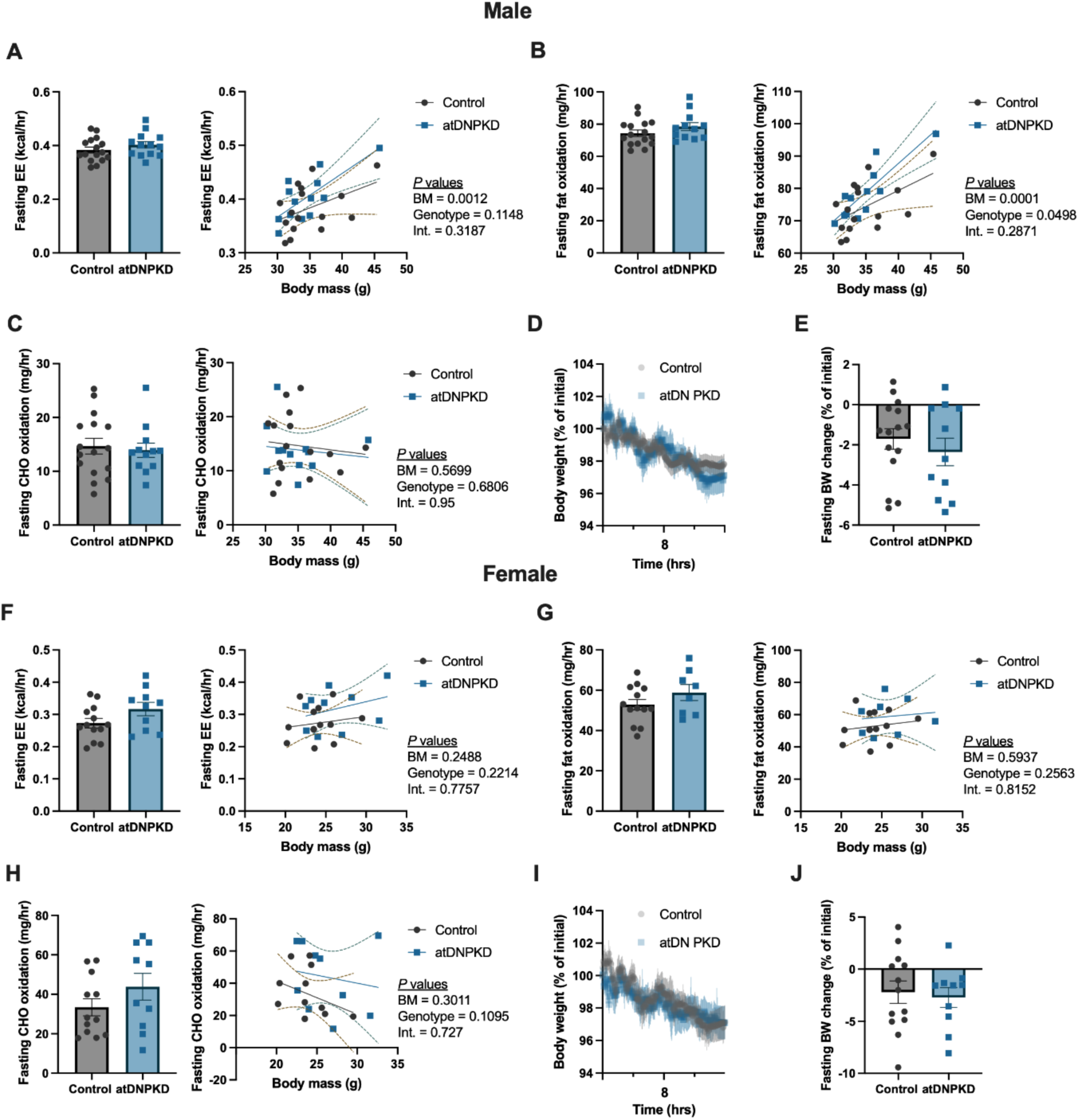
DNPKD expression in adipose tissue increases fat oxidation following an overnight fast in male (A-D), but not female (E-I) mice. Energy expenditure (EE; **A and F**), fat oxidation **(B and G)**, and carbohydrate (CHO) oxidation **(C and H)** in Control and atDNPKD mice during the final 4 hr of a 17 hr overnight fast. Body weight throughout the fasting period **(D and I)** and change in body weight (BW) due to fasting **(E and J)** in Control and atDNPKD mice. Data are mean ± SEM. n = 10-16/group. Displayed p-value obtained by independent samples t-test.

### Energy expenditure, substrate utilisation and feeding behaviour during refeeding in adipose tissue dominant negative PKD mice

A physiological situation in which it is necessary to restrain cAMP signalling in adipose tissue is during refeeding following fasting. As PKD activity is increased upon refeeding in adipose tissue (20), we hypothesised that adipose tissue PKD could have an important role in regulating metabolic responses during refeeding. Mice were housed in metabolic cages and following a 17 hr overnight fast, food was provided and metabolic responses over the next 4 hr were assessed. In male mice, there were no differences in energy expenditure (Figure 8A) or fat oxidation (Figure 8B) between groups. However, carbohydrate oxidation was lower in atDNPKD mice (Figure 8C). To understand whether the reduction in carbohydrate oxidation was associated with reduced exogenous carbohydrate availability, food intake over the first 4 hr of refeeding was assessed. Indeed, food intake was reduced in atDNPKD mice (Figure 8D) and remained lower 24 hr after refeeding (Figure 8E). To determine whether this impacted body weight, body weight regain following fasting was assessed (Figure 8F) and the time taken to regain the body weight lost due to fasting was longer in atDNPKD mice (Figure 8G). In female mice, there were no differences between groups in any of these parameters (Figure 8H-N). These data suggest that adipose tissue PKD plays a role in coordinating metabolic responses to fasting and refeeding that protect body weight in male mice.

**Figure 8:**
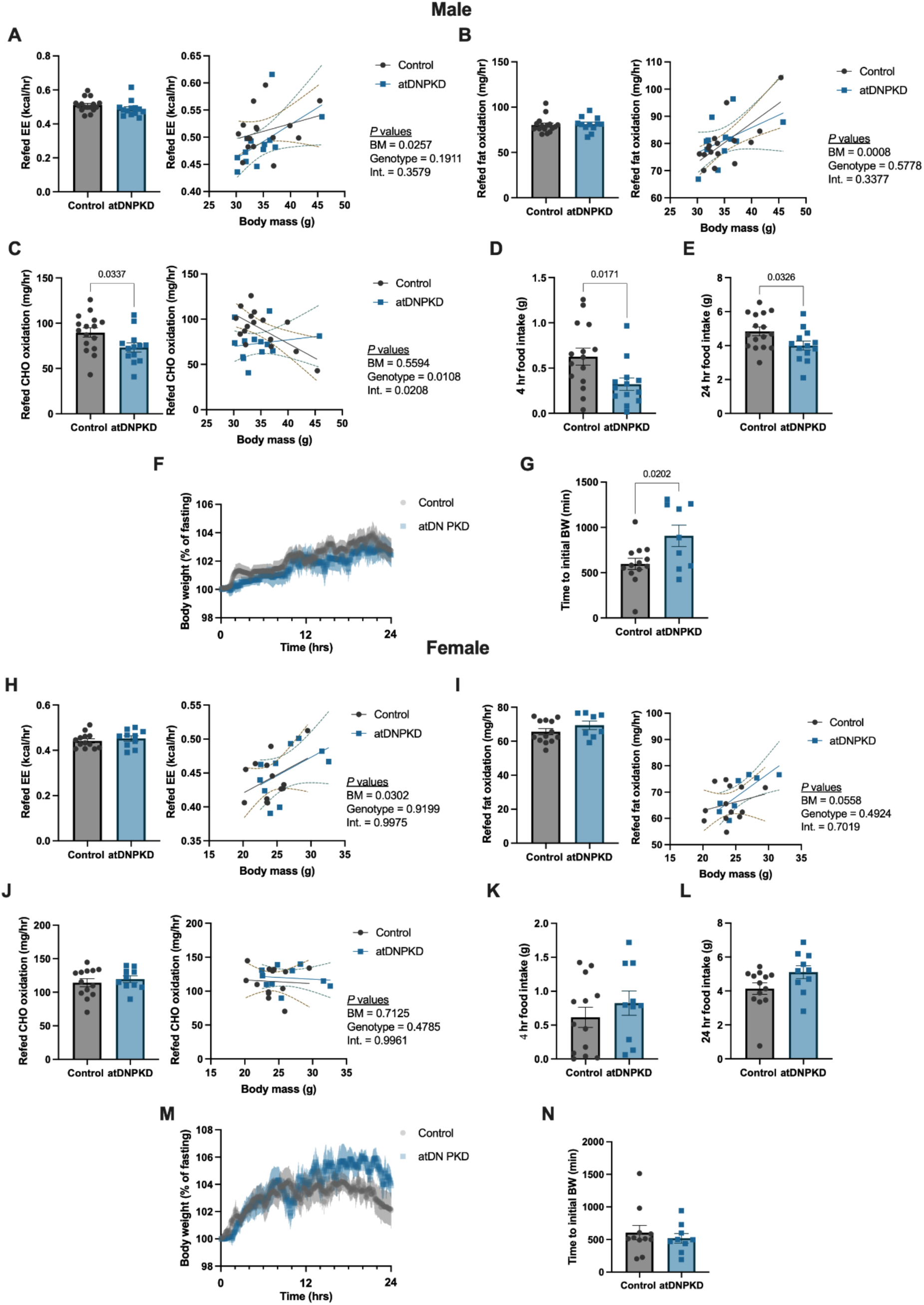
DNPKD expression in adipose tissue reduces food intake and increases time to body weight regain during refeeding following fasting in male (A-G), but not female (H-N) mice. Energy expenditure (EE; **A and H**), fat oxidation **(B and I)**, and carbohydrate (CHO) oxidation **(C and J)** during the first 4 hr of refeeding following a 17 hr overnight fast in Control and atDNPKD mice. Food intake in the first 4 hr **(D and K)** and 24 hr **(E and L)** of refeeding following a 17 hr overnight fast in Control and atDNPKD mice. Body weight throughout the refeeding period **(F and M)** and time to regain body weight (BW) lost due to fasting **(G and N)** in Control and atDNPKD mice. Data are mean ± SEM. n = 10-16/group. Displayed p-value obtained by independent samples t-test.

## DISCUSSION

Adipose tissue is the principal storage site for excess energy, and as such represents the frontline of obesity and related metabolic disorders. The PKD signalling pathway has become an exciting target for the development of novel therapeutics in the treatment for obesity and its comorbidities (11, 19). However, the physiological processes controlled by PKD and the downstream signalling pathways that mediate these effects remain largely unknown. The present study employed a phosphoproteomics approach to identify cellular signalling events regulated by PKD in differentiated 3T3-L1 mouse adipocytes. This approach revealed that PKD provided subtle regulation of pathways involved in metabolic control, including the cAMP and insulin signalling pathways, which reciprocally control key metabolic processes in adipose tissue. A novel inducible mouse model of adipocyte-specific inactivation of all three PKD isoforms (PKD1-3) was then employed to investigate the physiological and metabolic response to PKD inactivation *in vivo*. Results demonstrate that adipose tissue PKD plays a role in negative feedback control of cAMP signalling, which restrains energy expenditure and fat oxidation in response to β_3_-adrenoceptor activation. From a physiological standpoint, PKD plays a role in restraining fat oxidation during fasting, and food intake and body weight regain upon refeeding. These responses were observed in male but not female mice, highlighting important sex differences.

The PKD family of proteins are highly similar in structure leading to functional redundancy in multiple cell types (24–27). In the current study, this functional redundancy between isoforms was confirmed in differentiated 3T3-L1 adipocytes in culture. To effectively study the role of the PKD family in adipocytes while minimising functional redundancy, a siRNA-based *in vitro* triple PKD isoform knockdown model in 3T3-L1 differentiated adipocytes was developed. This approach, termed siPKD1/2/3, achieved an approximate 50% reduction in PKD family expression and phosphorylation.

With this siPKD1/2/3 adipocyte model, the signalling events downstream of PKD were investigated using the EasyPhos phosphoproteomics platform (30). As expected in a kinase knockdown model, the majority of PKD-regulated phosphosites were downregulated. Several phosphosites were upregulated but these are unlikely to be direct PKD substrates but rather involved in downstream pathways. Reductions in PKD isoform phosphorylation were observed across multiple unique phosphosites in response to PKD knockdown, confirming the knockdown observed via western blot sample validation. Of note, the most prominently expressed PKD phosphosite in PMA-treated 3T3-L1 cells, which was significantly reduced upon PKD knockdown, was PKD3 Ser^216^, a site conserved across both mouse and human but has not been addressed in the literature. Less is known regarding the activation process of PKD3 given it lacks the classical C-terminus autophosphorylation site of the other PKD isoforms (58). As such, the role of PKD3 Ser^216^ phosphorylation in activation and subsequent kinase action may be of interest in future research. No PKD1 phosphopeptides were detected by mass spectrometry using this workflow, however, gene and protein expression analysis does suggest that PKD1 expression was reduced in these samples.

Many of the identified differentially phosphorylated proteins were previously verified targets of PKD in other tissues. However, many hits have not previously been associated with PKD in any tissue. The most robust of these was transforming growth factor-β activated kinase 1 binding protein 2 (TAB2), which showed reduced PMA-stimulated phosphorylation at Ser^582^ with PKD knockdown. Importantly, this site is encased in a sequence motif highly homologous to previously identified PKD substrates, indicating that PKD might phosphorylate TAB2 directly. TAB2 is an adaptor protein central to the transforming growth factor-β activated kinase 1 (TAK1) inflammatory signalling cascade downstream of interleukin-1β (IL-1β) and tumour necrosis factor (TNF) (59). Given the inflammatory phenotype of adipose tissue in obesity, this stands out as a promising target for future research. However, the role of Ser^582^ phosphorylation on TAB2 function is currently unknown.

Analysis of differentially regulated phosphosites revealed PKD-mediated regulation of pathways involved in metabolism, including insulin and cAMP signalling that directly control adipose tissue glucose uptake, lipolysis, and energy expenditure. However, in differentiated 3T3L1 adipocytes, PKD1/2/3 knockdown had no effect on glucose uptake or lipolysis. Similarly, glucose kinetics were unchanged during an ITT in atDNPKD mice compared to control mice. Previous studies have demonstrated a role for PKD1 in cardiac glucose uptake (35, 51, 52), but this was contraction-induced glucose uptake, not insulin-stimulated. As such, PKD may not regulate insulin-stimulated glucose uptake, but it may be involved in other insulin-stimulated signalling pathways in adipocytes. For example, PKD activation in adipocytes has been linked to enhanced insulin-stimulated lipogenesis through inactivation of AMPK (20). Mitochondrial function was also assessed, which revealed an increase in mitochondrial uncoupling in PKD1/2/3 knockdown cells. This finding aligns with Loffler et al. (20) who showed increased mitochondrial fragmentation and uncoupled respiration in primary adipocytes that were isolated from adipose-specific PKD1 knockout mice. Similarly, RNA sequencing in visceral white adipose tissue from lean PKD1 knockout mice also found significantly upregulated UCP-1 gene expression when compared to control mice (21). This work strengthens the body of evidence suggesting that the PKD signalling pathway could be targeted to increase energy expenditure in adipose tissue in obesity.

To investigate adipose tissue PKD regulation of metabolism *in vivo*, an inducible, adipose-specific dominant negative PKD mouse model was developed which inhibits functional activity of all three PKD isoforms, an approach that has previously been used to study PKD function in the heart (35). Induction of the DNPKD transgene was observed in all three adipose depots, with the most robust induction in vWAT. Male atDNPKD mice had lower lean mass, but basal energy expenditure, and insulin-stimulated blood glucose kinetics were similar between atDNPKD and control mice for both sexes. These data are consistent with those of Loffler et al. (20), who found only minor improvements in insulin sensitivity in an insulin tolerance test in adipose tissue-specific PKD1 knockout mice, when compared with control mice. However, female atDNPKD mice displayed increased fasting plasma insulin and higher HOMA-IR index score. However, there were no differences in blood glucose between groups during an insulin tolerance test.

Given that adrenergic signalling and the cAMP signalling were identified in the phosphoproteomics screen as being regulated by PKD, the metabolic responses to β-adrenergic stimulation and fasting were further investigated. β-adrenergic stimulation increases lipolysis and adipokine release in adipocytes (57) and is critical for the thermogenic response in human brown/beige adipocytes (60). Results demonstrate that male, but not female, atDNPKD mice were more sensitive to β-adrenergic stimulation than control mice. Male atDNPKD mice showed modestly increased lipolysis, observed through an augmented increase in plasma FFA, and higher body weight-dependent energy expenditure and fat oxidation following β-adrenergic stimulation. These data suggest that PKD may fine-tune and limit the amplitude of β-adrenergic signalling in adipocytes in male mice. A multitude of phosphosites in the cAMP signalling pathway were found to be subtly regulated by PKD which also suggests dampening of this pathway. One context in which this regulatory mechanism could be relevant is fasting, where PKD activation in adipose tissue could act to balance substrate demand and energy expenditure to preserve body weight. This hypothesis was tested by studying mice during an overnight fast, and although fasting fat oxidation appears to be controlled by PKD, no differences in energy expenditure or body weight were observed between groups. Whether more extreme forms of metabolic perturbation, such as extended cold exposure or fasting, might identify the evolutionary role of PKD in adipose tissue remains to be determined. However, an important physiological role for adipose PKD activity during refeeding following a period of fasting was observed, where adipocyte PKD is important for promoting food intake and restoring body weight. The mechanisms involved remain to be determined, however one potential mechanism could be related to the effects of adiponectin on food intake. Inhibition of PKD increases adiponectin secretion from cultured adipocytes (28) and adiponectin has inhibitory effects on food intake after fasting (61). The mechanism uncovered in this study appears to be specific for fasting and refeeding, as there were no differences in indices of energy balance between the groups of mice under free living conditions where food was provided *ad libitum* (Figure 4).

Given the demonstrated redundancy in PKD isoforms in the current study and from others (24–27), inhibition of all three PKD isoforms is a major strength of the models used in this study. However, results in the 3T3-L1 adipocytes only indicate an approximate 50% reduction in PKD phosphorylation. This highlights the trade-off between partial knockdown of all three PKD isoforms, and achieving complete knockdown in one isoform alone, as has been employed previously (20, 21, 24, 28). Future research will need to establish both unique functions of individual isoforms, and redundant functions regulated by all three isoforms. The relatively low number of significantly differentially regulated phosphosites following multiple comparison testing (26 were identified) could be a consequence of the incomplete PKD knockdown in the adipocyte model. As such, the 432 differentially regulated phosphosites before multiple comparison testing were used for pathway analysis, although a limitation to this approach is the increased likelihood of multiple type I errors.

Interrogating the 3T3L1 phosphoproteome in this study generated rich data on the protein activation patterns downstream of PKD. However, signalling pathways are also regulated by other post-translational modifications, and by the total expression of key proteins. Therefore, future research should attempt to measure the total proteome alongside the phosphoproteome following PKD inactivation in adipocytes. Multiomics approaches such as this can provide a more complete picture of cellular signalling pathways in adipocytes.

This study highlighted important sex-differences in the regulation of adipose tissue metabolism by PKD. However, the lack of effect of atDNPKD expression in female mice compared with male mice is challenging to reconcile. It has been noted that low dose tamoxifen exposure is sufficient to induce persistent metabolic changes in female mice (62). As such, it is possible that tamoxifen-induced metabolic adaptations were sufficient to mask the subtle effects of PKD on metabolic control and an alternate inducible Cre model could be required to properly assess the functional role of adipose PKD in female mice.

In conclusion, results suggest that PKD plays a regulatory role in fine tuning metabolic pathways, particularly the cAMP and insulin signalling pathways in adipose tissue. From a functional standpoint, PKD likely tempers the magnitude of adrenergic signalling on adipose tissue in male, but not female, mice which has functional consequences on metabolic and behavioural responses under conditions such as fasting and refeeding.

## GRANTS

This work was supported by the National Health and Medical Research Council (APP1163238 to SLM).

## AUTHOR CONTRIBUTIONS

M.C.R., S.J.H., D.E.J., K.F.H., and S.L.M. conceived and designed research; M.C.R., S.J.H., T.C. S.D.M., K.K., H.F. K.F.H., S.L.M. performed experiments; M.C.R., S.J.H., T.C. S.D.M., K.K., H.F. analysed data; M.C.R., S.J.H., D.E.J., K.F.H., and S.L.M. interpreted results of experiments; M.C.R., S.J.H., K.F.H., and S.L.M. prepared figures; M.C.R., S.J.H., K.F.H., and S.L.M drafted manuscript; M.C.R., S.J.H., C.S.S., K.F.H., and S.L.M. edited and revised manuscript; All authors approved final version of manuscript.

