## Supplemental Material for "Adipose tissue protein kinase D (PKD): regulation of signalling networks and its sex-dependent effects on metabolism"

**RUNNING HEAD:** PKD regulates adipose tissue metabolism

**CORRESPONDING AUTHORS:**

Kirsten F. Howlett
Sean L. McGee

### SUPPLEMENTARY FIGURE 1

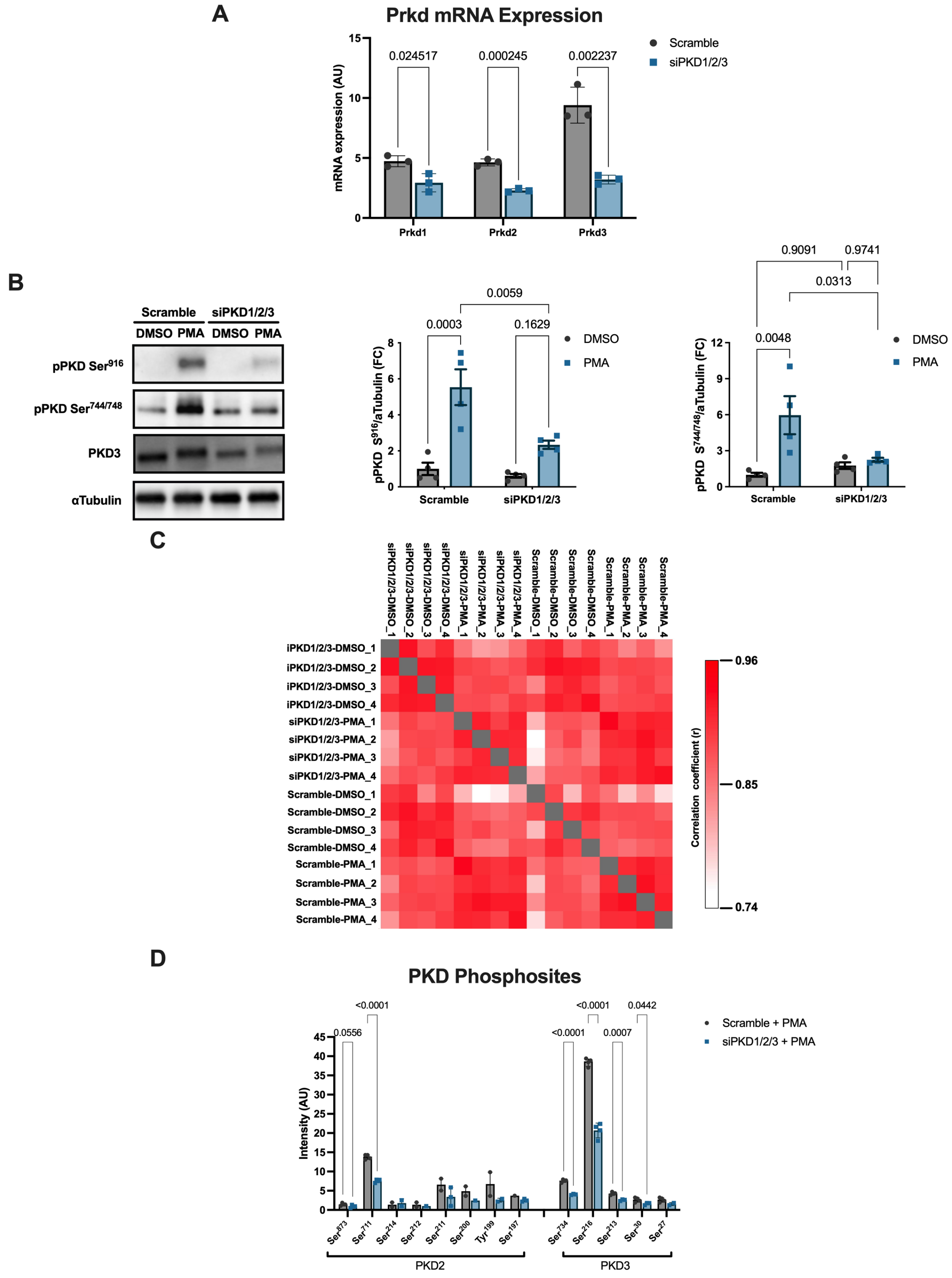

**Supplementary Figure 1: Phosphoproteomics sample validation.**

A. *Prkd* mRNA expression analysis via real time qPCR in scramble and siPKD1/2/3 transfected differentiated 3T3-L1 adipocytes.
B. Phosphorylated PKD protein expression via western blot in scramble and siPKD1/2/3 transfected adipocytes treated with either PMA or DMSO vehicle for 20 min. Corrected for $\alpha$Tubulin expression as protein loading.
C. Pearson's correlation analysis of individual phosphoproteomic samples.
D. Expression of PKD family phosphopeptides compared between PMA treated scramble and siPKD1/2/3 adipocytes.

**SUPPLEMENTARY FIGURE 2**

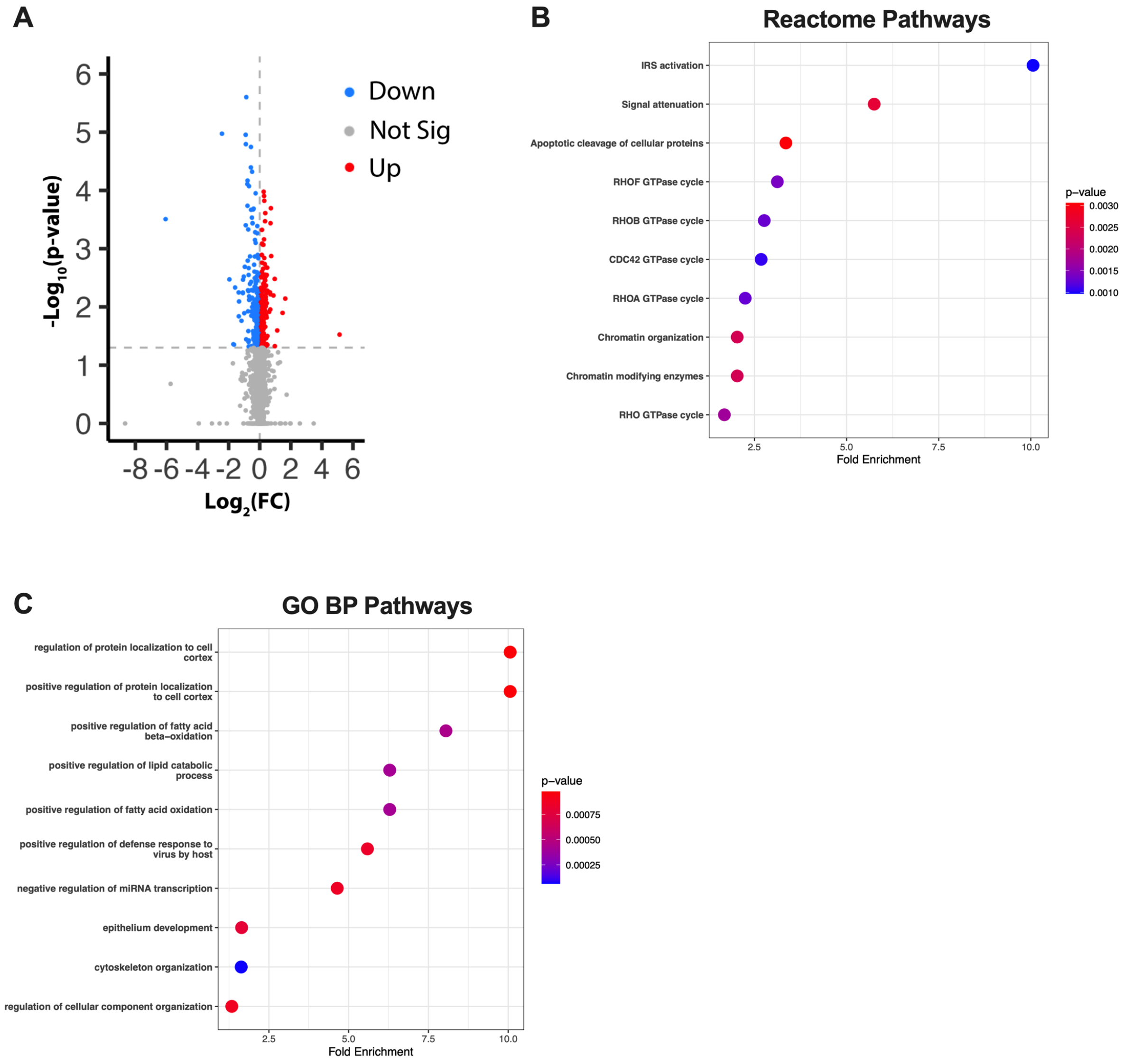


**Supplementary Figure 2: Pathway analysis of identified phosphosites downstream of PKD in differentiated 3T3-L1 adipocytes.**

A. Volcano plot displaying the -$Log_{10}$ p-value from independent samples t-test and the $Log_2$ fold change in expression of phosphosites between PMA treated scramble and siPKD1/2/3 adipocytes.
B. Top ten overrepresented Reactome pathways by unadjusted p-value significance ranked by fold change identified in the 432 differentially regulated phosphosites before multiple comparison testing.
C. Top ten overrepresented Gene Ontology Biological Processes (GO BP) pathways by unadjusted p-value significance ranked by fold change identified in the 432 differentially regulated phosphosites before multiple comparison testing.

SUPPLEMENTARY FIGURE 3

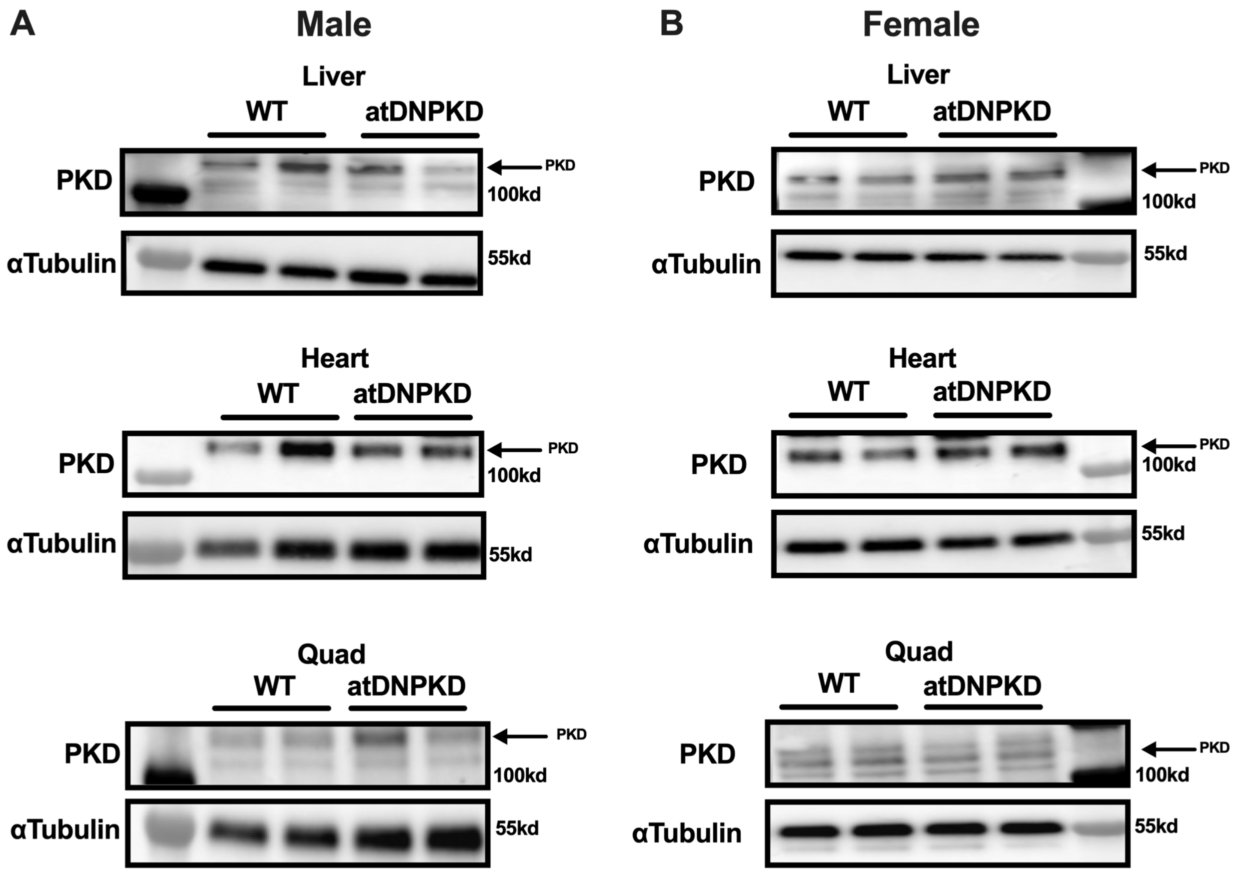


**Supplementary Figure 3: DNPKD transgene expression is specific to adipose tissue.**

A. PKD protein expression via western blot in liver, heart, and skeletal muscle (quad) protein lysates from male WT and atDNPKD mice. $\alpha$Tubulin displayed as loading control. n = 2 per group.

B. PKD protein expression via western blot in liver, heart, and skeletal muscle (quad) protein lysates from female WT and atDNPKD mice. $\alpha$Tubulin displayed as loading control. n = 2 per group.
